# *Mycobacterium tuberculosis* requires riboflavin biosynthesis for survival in the host

**DOI:** 10.64898/2026.09.25.754416

**Authors:** Alisha M. Block, Sarah B. Namugenyi, Anna D. Tischler

**Author notes:** Corresponding author. (ADT). Sequencing and Bioinformatics Unit, Minnesota Department of Health, St. Paul, MN, USA.

## Abstract

*Mycobacterium tuberculosis* has a flavin-intensive metabolism, with more than 180 proteins that require flavin or deazaflavin cofactors. *M. tuberculosis* requires riboflavin biosynthesis for growth in standard culture conditions *in vitro*, which can be overcome by providing exogenous riboflavin. It is unknown whether *M. tuberculosis* also requires riboflavin synthesis to grow or persist in infected host cells or tissues. Here we used a *M. tuberculosis* strain that conditionally expresses *ribA2*, which encodes a bifunctional enzyme that catalyzes the first steps in both branches of riboflavin biosynthesis, to determine whether *M. tuberculosis* requires riboflavin biosynthesis during infection. Transcriptional repression of *ribA2* caused death of *M. tuberculosis in vitro* without riboflavin supplementation. RNAseq revealed that riboflavin starved *M. tuberculosis* induce a transcriptional program indicating decreased electron transport chain function and broad changes to central metabolism. Transcriptional repression of *ribA2* prevented *M. tuberculosis* growth in cultured macrophages and caused clearance of *M. tuberculosis* from infected tissues at all stages of infection in both C57BL/6 mice that form cellular lung lesions and C3HeB/FeJ mice that develop necrotic lung granulomas. Our results demonstrate that *M. tuberculosis* requires RibA2 for viability at all stages of infection and support further characterization of the riboflavin biosynthesis pathway as a potential target for development of new anti-tubercular drugs.

**AUTHOR SUMMARY:** Tuberculosis (TB) caused by *Mycobacterium tuberculosis* kills over 1.2 million people worldwide each year, making it the deadliest bacterial infection. TB treatment involves combinations of antibiotics taken for at least six months. *M. tuberculosis* readily becomes resistant to antibiotics by mutation and multidrug resistant strains account for nearly 4% of new TB cases. New antibiotics for TB are needed to combat drug resistance and shorten TB treatment. Riboflavin biosynthesis is an attractive target for antibiotic development because mammals do not synthesize riboflavin, instead obtaining it from the diet. Here, using a *M. tuberculosis* strain that conditionally expresses the riboflavin biosynthesis enzyme RibA2, we show that inhibition of riboflavin biosynthesis kills *M. tuberculosis* in lab culture conditions and in mice at all stages of infection. Our results demonstrate that riboflavin starvation is bactericidal for *M. tuberculosis* and identify the riboflavin biosynthesis pathway as a promising target for new drugs to treat TB.

## INTRODUCTION

New antibiotics with unique molecular targets are needed to treat tuberculosis (TB) caused by *Mycobacterium tuberculosis*, which killed more people worldwide in 2024 than any other infectious agent (1). Treatment of both drug-susceptible and drug-resistant *M. tuberculosis* infections requires at least 6 months of combination therapy with multiple antibiotics (2, 3). *M. tuberculosis* readily evolves drug resistance by mutation, which can lead to treatment failure (4–6). New antibiotic regimens for multidrug resistant TB have improved treatment outcomes (7, 8). However, emergence of *M. tuberculosis* strains that are resistant to these new drugs and that are readily transmitted threatens the continued efficacy of these drug regimens (9–11). There is therefore an urgent need to identify new targets for development of novel anti-tubercular drugs that can be used in combination therapy with existing antibiotics to both shorten TB treatment and combat emerging drug resistance.

Central metabolic pathways that *M. tuberculosis* requires for survival in the host and that are absent in humans are potential targets for anti-tubercular drug development. Genome-wide transposon (Tn) mutagenesis screens identified genes that *M. tuberculosis* requires to grow in lab culture conditions (12, 13) and CRISPR interference (CRISPRi) further defined the vulnerability of these pathways to inhibition (14). Based on these screens, many central metabolic pathways are equally vulnerable to inhibition as the targets of existing antibiotics (14). We previously screened a small collection of auxotrophic *M. tuberculosis* mutants lacking central metabolic functions in intravenously infected mice and identified multiple pathways that were essential for survival in the host including biosynthesis of purine nucleotides, amino acids (Phe, Trp, Pro, Glu), and riboflavin (15). *M. tuberculosis* mutants defective for riboflavin biosynthesis (*ribA2*::Tn, *ribG*::Tn) were among the most strongly attenuated in our screen (15). Mammals do not encode the enzymes for *de novo* riboflavin biosynthesis and must instead obtain riboflavin from the diet (16), suggesting that antibiotics targeting riboflavin biosynthesis would have limited toxicity. Riboflavin biosynthesis is therefore a promising target for anti-tubercular drug development.

Flavin cofactors produced from riboflavin play many essential physiological functions in *M. tuberculosis*. Riboflavin is a precursor for the flavin cofactors flavin mononucleotide (FMN) and flavin adenine dinucleotide (FAD). Riboflavin biosynthesis intermediates are also used to produce the F_420_ deazaflavin cofactor. *M. tuberculosis* metabolism is highly flavin dependent, with an estimated 150 flavoproteins that use FMN or FAD and 33 F_420_-dependent proteins (17, 18). FMN and FAD are versatile redox cofactors capable of performing both one- and two-electron transfer reactions in enzymes with diverse catalytic mechanisms (19). *M. tuberculosis* requires flavin cofactors for components of the electron transport chain that enable respiratory metabolism, enzymes involved in metabolism of carbon sources including fatty acids and cholesterol, and redox homeostasis (16). Although *M. tuberculosis* does not require the F_420_ co-factor for viability (12), mutants lacking F_420_ are highly susceptible to oxidative and nitrosative stress and certain antibiotics (20, 21). Thus, inhibition of riboflavin biosynthesis is expected to have pleiotropic effects on *M. tuberculosis* physiology.

The *M. tuberculosis* riboflavin biosynthesis pathway includes multiple vulnerable enzymatic steps that are potential targets for drug development. Riboflavin is biosynthesized from ribulose-5-phosphate derived from the pentose phosphate pathway and GTP precursors in a process requiring five enzymes: RibA2, RibG, RibC, RibH and an unidentified phosphatase (16). Riboflavin is then converted to FMN and FAD by RibF (16). The F_420_ deazaflavin cofactor is produced from the riboflavin biosynthesis intermediate 5-amino-6-D-ribitylaminoaminouracil (5-A-RU) by FbiA-D (16). Conditional silencing of *ribA2*, *ribG*, or *ribH* transcription using CRISPRi limited *M. tuberculosis* growth *in vitro*, demonstrating the vulnerability of this pathway to inhibition (22, 23). RibA2 is a bi-functional enzyme required for the rate-limiting first steps of riboflavin synthesis from both ribulose-5-phosphate and GTP (24). Genome-wide CRISPRi screens identified *ribA2* as a highly vulnerable target in standard culture media *in vitro*, in macrophages, and in an *ex vivo* mouse tissue culture infection model (14, 25, 26). However, it is unknown whether inhibiting riboflavin biosynthesis kills *M. tuberculosis* in host tissues during acute or chronic infection.

Here we use conditional expression of RibA2 to show that *M. tuberculosis* requires riboflavin biosynthesis for survival *in vitro* without riboflavin supplementation and at all stages of infection in mice. RibA2 depletion and starvation for riboflavin trigger a transcriptional program indicating reduced electron transport chain function and altered central carbon and fatty acid metabolism. Exogenous riboflavin partially rescued growth of RibA2-deficient *M. tuberculosis* in a macrophage infection model. However, there is insufficient riboflavin in host tissues to rescue growth of RibA2-deficient *M. tuberculosis*, as transcriptional repression of *ribA2* caused rapid clearance of *M. tuberculosis* from infected tissues at all stages of infection in C57BL/6 mice. *M. tuberculosis* also required RibA2 for persistence in C3HeB/FeJ mice that develop necrotic granulomas. Our results support the riboflavin biosynthesis pathway, and RibA2 specifically, as a potential target for development of new anti-tubercular drugs.

## RESULTS

### *M. tuberculosis ribA2*::Tn cannot grow without riboflavin supplementation

In our Tn-seq screen, we showed that a *M. tuberculosis ribA2*::Tn mutant was highly attenuated in intravenously infected mice (15). The *ribA2*::Tn mutant could not grow in standard Middlebrook 7H9 medium but grew normally in MtbYM rich medium that contains 53 μM riboflavin (15). To determine if riboflavin starvation is cidal, we examined survival of *ribA2*::Tn and *ribC*::Tn mutants isolated from our Tn mutant library (27) in 7H9 medium with and without 53 μM riboflavin. The *ribA2*::Tn and *ribC*::Tn mutants failed to grow and slowly lost viability in 7H9 without riboflavin but grew similarly to the WT control and remained viable with 53 μM riboflavin (**Fig 1A-B**). These data confirm that *M. tuberculosis* requires RibA2 and RibC for viability in standard culture conditions *in vitro*.

**Figure 1.**
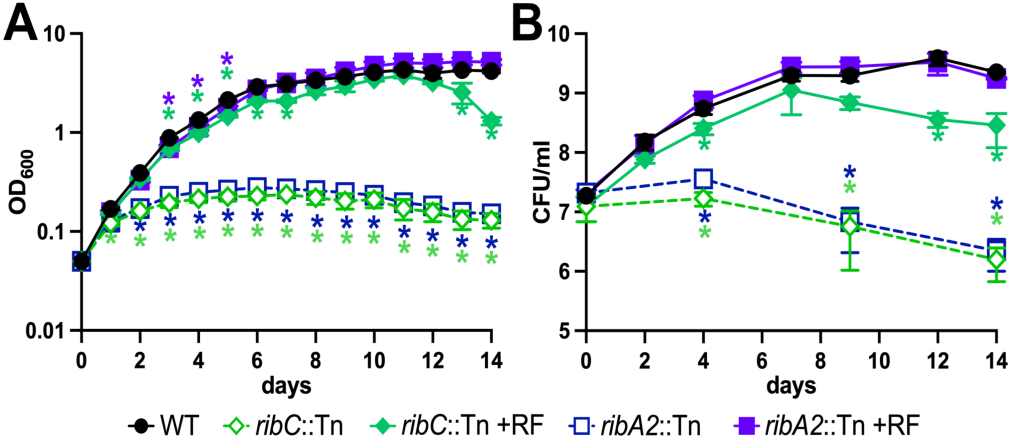
*M. tuberculosis ribA2*::Tn and *ribC*::Tn mutants lose viability without exogenous riboflavin. WT Erdman, *ribA2*::Tn and *ribC*::Tn were grown in complete 7H9 (WT) or complete 7H9 with 53 μM riboflavin (7H9+RF; Tn mutants) to mid-log phase, washed and diluted to an OD_600_ of 0.01 in complete 7H9 or 7H9+RF. Growth was monitored by OD_600_ measurement (**A**) and plating serially diluted cultures on 7H10 agar (WT) or on 7H10 agar with 53 μM riboflavin (Tn mutants) (**B**). Data are means ± standard deviations of triplicate cultures. Asterisks indicate significant differences between the Tn mutant and the WT control by one-way ANOVA with Dunnett’s multiple comparisons test post-hoc: \**P*<0.05.

### Transcriptional repression of *ribA2* causes death of *M. tuberculosis* without exogenous riboflavin

To evaluate the requirement for RibA2 during *M. tuberculosis* infection, we generated a *ribA2* Tet-OFF conditional expression strain. We cloned *ribA2* under the control of a tetracycline-repressible promoter on an integrating plasmid and then deleted the native copy of *ribA2* (**Fig S1A-B**). Whole-genome sequencing of *ribA2* Tet-OFF did not identify any secondary mutations in the locus required for production of phthiocerol dimycocerosate (PDIM), an outer mycolate layer lipid required for *M. tuberculosis* virulence (28). The *ribA2* Tet-OFF strain produces PDIM (**Fig S1C**). The *ribA2* Tet-OFF strain grows without riboflavin but at a slower rate than the WT control (**Fig 2A-B**). This slower growth rate was not rescued by exogenous riboflavin (**Fig 2C-D**). Treatment of *ribA2* Tet-OFF with anhydrotetracycline (Atc) repressed *ribA2* transcription 7-fold (**Fig S1D**). Expression of *ribH*, which is encoded in an operon with *ribA2* (**Fig S1A**), was also 6-fold lower in *ribA2* Tet-OFF relative to the WT control, even without Atc (**Fig S1E**), possibly due to the deletion of a transcription start site within *ribA2* (29). Decreased expression of *ribH* may contribute to the growth defect of *ribA2* Tet-OFF in the absence of riboflavin.

**Figure 2.**
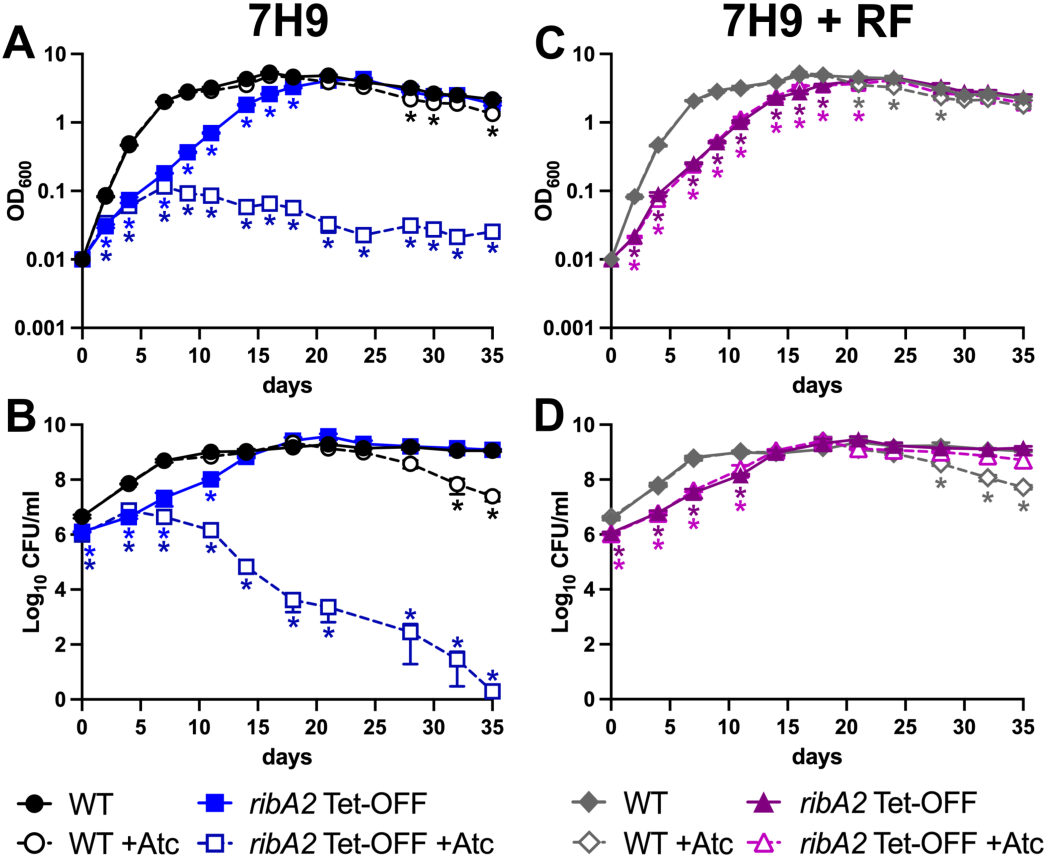
*M. tuberculosis* requires RibA2 to survive in the absence of exogenous riboflavin. WT Erdman and *ribA2* Tet-OFF were grown in complete 7H9 medium to mid-log phase, washed and diluted to an OD_600_ of 0.01 in complete 7H9 (**A, B**) or complete 7H9 with 53 μM riboflavin (7H9+RF; **C, D**) ± 100 ng/ml anhydrotetracycline (Atc). Growth was monitored by OD_600_ measurement (**A, C**) and by plating serially diluted cultures on 7H10 agar (WT) or on 7H10 with 53 μM riboflavin (*ribA2* Tet-OFF) (**B, D**). Data are means ± standard deviations of triplicate cultures. Asterisks indicate significant differences from the WT by one-way ANOVA with Dunnett’s multiple comparisons test post-hoc. \**P*<0.05.

To determine if riboflavin biosynthesis inhibition is cidal without exogenous riboflavin, we grew *ribA2* Tet-OFF in 7H9 medium with Atc ± 53 μM riboflavin. Atc prevented growth of *ribA2* Tet-OFF, but this growth defect was rescued with exogenous riboflavin (**Fig 2A&C**). To quantify *ribA2* Tet-OFF viable CFU, we plated cultures on 7H10 medium supplemented with 53 μM riboflavin. Transcriptional repression of *ribA2* with Atc caused rapid death of *M. tuberculosis*, with near sterilization of the cultures in 35 days (**Fig 2B**). Exogenous riboflavin prevented death of the *ribA2* Tet-OFF strain with Atc (**Fig 2D**). These data show that *M. tuberculosis* requires riboflavin biosynthesis for viability in standard culture conditions *in vitro*, but that this requirement can be overcome with exogenous riboflavin.

### Repression of *ribA2* alters expression of respiration and central carbon metabolism genes

To determine the physiological effect of riboflavin starvation, we analyzed the transcriptional profile associated with *ribA2* repression by RNAseq. We grew WT Erdman or *ribA2* Tet-OFF in 7H9 medium without riboflavin ± Atc. For WT, we included an ethanol solvent control. We collected bacteria for RNA extraction at day 1 for WT and days 1, 2 and 4 for *ribA2* Tet-OFF. In growth curves, we did not observe significant loss of viability of *ribA2* Tet-OFF until after 4 days of Atc treatment (**Fig 2B**). We compared all transcriptomes to the WT no Atc control (**Table S1**). We identified only a few genes that were differentially expressed by WT *M. tuberculosis* grown either +Atc (**Fig S2A**) or +ethanol (**Fig S2B**), suggesting limited impact of these conditions on gene expression.

The *ribA2* Tet-OFF no Atc control exhibited significantly increased expression of 105 genes and decreased expression of 53 genes at day 1 (**Fig 3A, Table S1**). These include up-regulation of *prpDC* and *icl1*, which encode methylcitrate cycle enzymes involved in propionate catabolism (30) (**Figs 3A & 3D**), and down-regulation of genes in the *mce1* operon that is required for fatty acid uptake (31) and phospholipases *plcABC* involved in fatty acid utilization (32) (**Fig S3A**). These genes were also differentially expressed by *ribA2* Tet-OFF relative to WT at days 2 and 4 and were more strongly induced or repressed by Atc treatment (**Figs 3D & 3E, Fig S3A & S3B**). These data indicate that propionate and lipid metabolism are already disrupted in the *ribA2* Tet-OFF strain, even without transcriptional repression of *ribA2*.

**Figure 3.**
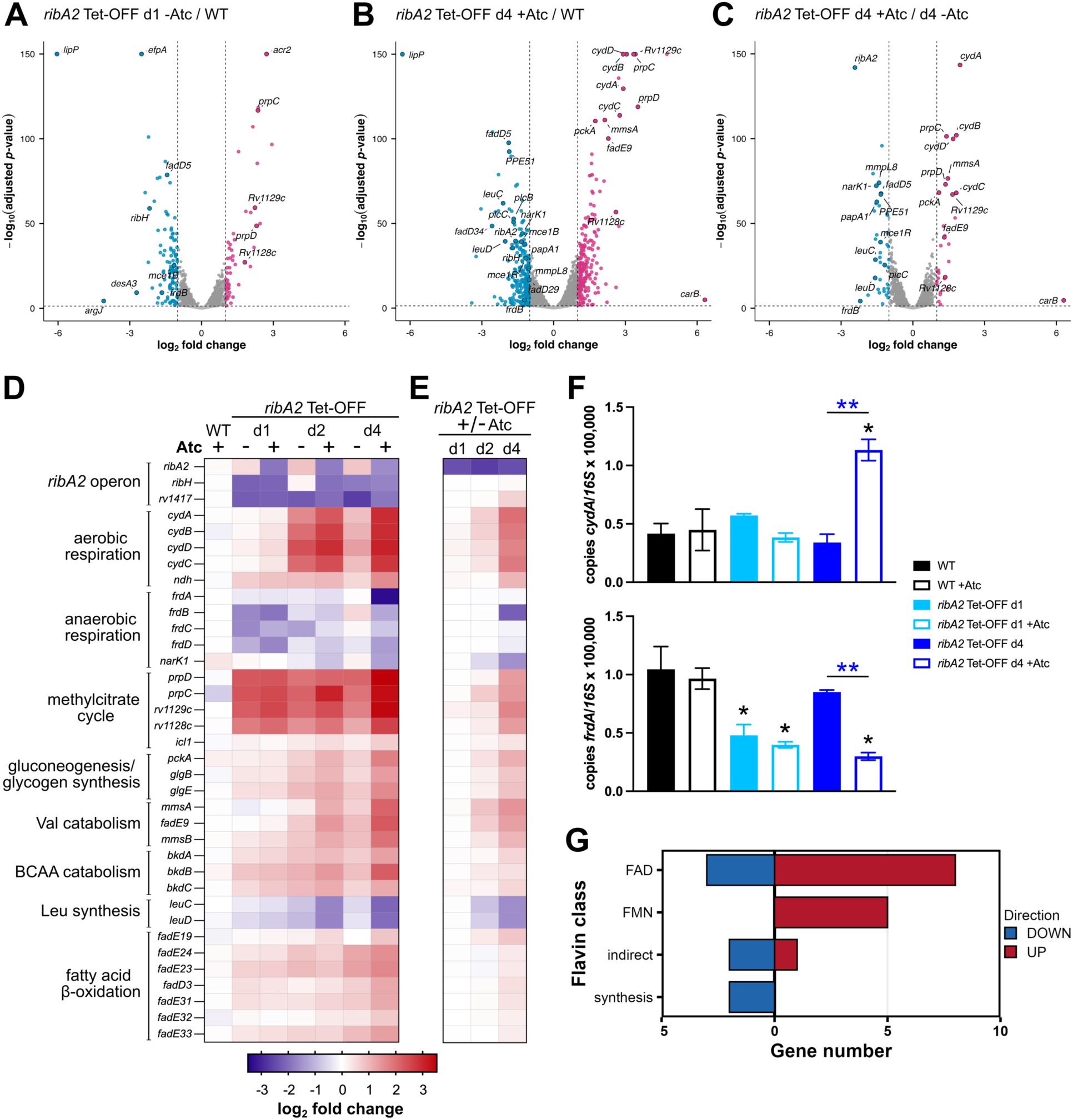
Transcriptional repression of *ribA2* alters expression of genes encoding electron transport chain components and central metabolic processes. RNA was extracted from WT Erdman or *ribA2* Tet-OFF grown in 7H9 ± 100 ng/ml Atc and transcriptional profiles were determined by RNAseq. **A-C.** Volcano plots comparing transcript abundance in *ribA2* Tet-OFF no Atc at day 1 to the WT no Atc control (**A**), *ribA2* Tet-OFF +Atc at day 4 to the WT no Atc control (**B**), or *ribA2* Tet-OFF +Atc at day 4 to the *ribA2* Tet-OFF no Atc control at day 4 (**C**). Each dot represents an individual gene. Dashed lines indicate the log_2_ fold change = ± 1 and *Padj* < 0.05 statistical significance cutoffs. Red or blue dots represent significantly up- or down-regulated genes relative to the control, respectively. **D-E.** Heat maps showing the log_2_ fold change in transcript abundance of select differentially expressed genes across all samples relative to the WT no Atc control (**D**) or relative to the *ribA2* Tet-OFF no Atc control from the same time point (**E**). **F.** Abundance of the *cydA* and *frdA* transcripts relative to 16S rRNA determined by qRT-PCR. Data are means ± standard deviations of biological triplicate samples. Black asterisks indicate statistically significant differences from the WT no Atc control by one-way ANOVA with Dunnett’s multiple comparisons test post-hoc. Blue asterisks indicate statistically significant differences ± Atc by unpaired *t*-test. \**P*<0.05, \*\**P*<0.001. **G.** Number of significantly up- or down-regulated genes encoding flavin-related proteins in *ribA2* Tet-OFF +Atc day 4 compared to the WT no Atc control.

Addition of Atc caused limited changes to the *ribA2* Tet-OFF transcriptome at days 1 and 2 relative to either the WT control (**Fig 2D**) or the no Atc control from the same time point (**Fig 2E**). Only transcription of *ribA2* itself was significantly reduced in these comparisons (**Fig S2C & S2D**). These results are consistent with our observation that *ribA2* transcriptional repression did not cause loss of viability until after 4 days of culture without riboflavin (**Fig 2B**).

At day 4, we observed significant changes in gene expression in *ribA2* Tet-OFF grown +Atc relative to both the WT control (**Fig 3B**) and the *ribA2* Tet-OFF no Atc control (**Fig 3C**). The *cydABDC* operon encoding the proton pumping cytochrome *bd* terminal oxidase (Cyt-*bd*) was strongly upregulated (**Fig 3B-E**). Cyt-*bd* is important for electron transport chain function and respiration in hypoxic conditions (33). The *M. tuberculosis* NADH dehydrogenases NDH-1 (*nuoA-N*) and NDH-2 (*ndh* and *ndhA*), which serve as entry points for transfer of electrons from NADH into the respiratory chain, require the FMN and FAD cofactors, respectively (34). *M. tuberculosis* may increase Cyt-*bd* expression to compensate for decreased proton pumping by NDH-1 when riboflavin biosynthesis is inhibited. We also observed up-regulation of *ndh* encoding NDH-2 and down-regulation of *frdABCD* and *narK1* (**Fig 3B-E**), which encode fumarate reductase and a nitrate/nitrite transporter, respectively. Fumarate and nitrate can serve as alternative terminal electron acceptors in anaerobic conditions (34). We validated increased expression of *cydA* and decreased expression of *frdA* in *ribA2* Tet-OFF grown +Atc (**Fig 3F**). Our data suggest that *ribA2* transcriptional repression alters electron transport chain activity.

Transcriptional profiling also revealed decreased expression of genes required for Leu biosynthesis (*leuCD*) and increased expression of genes required for branched-chain amino acid catabolism (*mmsA*, *fadE9*, *mmsB*, *bkdABC*), gluconeogenesis and glycogen synthesis (*pckA*, *glgB*, *glgE*), and fatty acid β-oxidation (*fadE* genes) upon *ribA2* repression with Atc (**Fig 3B-E**). These changes in expression of genes encoding central metabolic functions are likely due to decreased activity of enzymes that require a flavin cofactor. Indeed, we identified many differentially expressed genes that encode enzymes either directly or indirectly associated with flavin cofactors (**Fig 3G, Fig S3C**). These include BkdB, a component of the branched-chain keto acid dehydrogenase complex, and all FadE enzymes that require FAD (35, 36). Genes encoding these enzymes may be up-regulated to compensate for decreased enzyme activity when riboflavin biosynthesis is inhibited.

Other genes may be differentially expressed due to indirect effects on flavin-dependent metabolic processes. Lipoamide dehydrogenase (Lpd), a component of pyruvate dehydrogenase that converts pyruvate to acetyl-CoA, requires FAD (36, 37). Decreased Lpd activity is expected to increase availability of pyruvate. *M. tuberculosis* growth on pyruvate depends on a reverse methyl citrate cycle (PrpDC, Icl1), the gluconeogenic enzyme PckA, and MmsA, which is involved in Val catabolism (38). Genes encoding these enzymes were all up-regulated upon *ribA2* repression (**Fig 3B-E**), suggesting a compensatory response to increased intracellular pyruvate. MmsA and MmsB may also be upregulated to enable NAD recycling, as both enzymes catalyze reversible reactions that can use NADH to produce NAD^+^ (39, 40). Regeneration of NAD^+^ is expected to be disrupted in cells depleted of flavins because the flavin-dependent NADH dehydrogenases NDH-1 and NDH-2 are the primary route for NAD recycling (34, 41). Cyt-*bd* does not use NADH, so it cannot compensate for this function. We also observed decreased expression of genes required for lipid synthesis and genes encoding ribosomal proteins (**Fig S3A & S3B**), consistent with slower growth upon *ribA2* repression. Our data indicate rerouting of carbon through central metabolism, lipid utilization and lipid synthesis pathways when riboflavin synthesis is inhibited.

### *M. tuberculosis* requires RibA2 to grow in resting and activated macrophages

To determine the importance of riboflavin biosynthesis to *M. tuberculosis* growth in infected cells, we used the human THP-1 monocyte cell line differentiated to macrophages. THP-1 cells were infected with WT Erdman or *ribA2* Tet-OFF and cultured ± Atc. Repression of *ribA2* with Atc was initiated at either the start of the THP-1 infection (day 0) or during growth of the pre-culture used for infection (pre). Atc had no effect on growth of WT *M. tuberculosis* in THP-1 cells (**Fig 4A**). The *ribA2* Tet-OFF strain grew normally in resting THP-1 cells, but repression of *ribA2* with Atc prevented its replication (**Fig 4A**). Growth inhibition was most effective with Atc treatment of the pre-culture (**Fig 4A**). Activation of THP-1 cells with IFN-γ did not improve control of WT Erdman, but partially restricted growth of the *ribA2* Tet-OFF strain, even without Atc (**Fig 4B**). IFN-γ activated THP-1 cells exhibited slightly improved control of the *ribA2* Tet-OFF strain when *ribA2* was repressed with Atc (**Fig 4B**). These data indicate that *M. tuberculosis* cannot acquire sufficient riboflavin from resting or IFN-γ activated THP-1 cells to support its growth.

**Figure 4.**
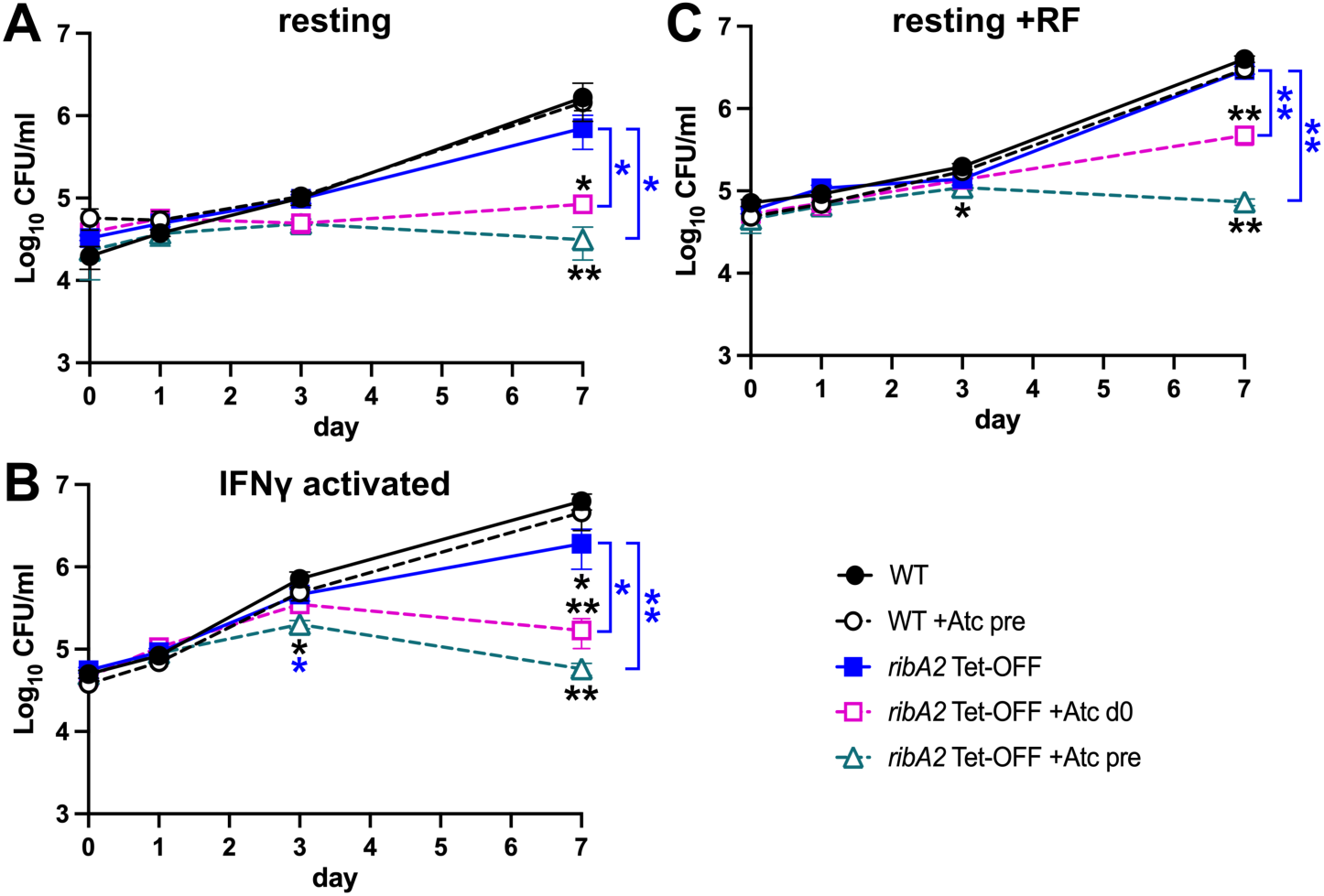
*M. tuberculosis* requires RibA2 for growth in macrophages. Resting (**A, C**) or interferon-gamma (IFN-γ) activated (**B**) THP-1 cells were infected at a MOI of 1:20 bacteria:macrophage with WT Erdman or *ribA2* Tet-OFF grown to mid-log phase in 7H9 or 7H9 + 100 ng/ml anhydrotetracycline (Atc) (pre). Macrophages were cultured ± 200 ng/ml Atc. In **C**, macrophages were cultured with 53 μM riboflavin. Viable intracellular CFU were determined by plating cell lysates on 7H10 agar (WT) or 7H10 with 53 μM riboflavin (*ribA2* Tet-OFF). Data are means ± standard deviations of duplicate wells from at least three biological replicates. Asterisks indicate statistically significant differences from the WT (black) or the *ribA2* Tet-OFF no Atc control (blue) by one-way ANOVA with Dunnett’s multiple comparisons test post-hoc: \**P*<0.05, \*\**P*<0.001.

To determine if the growth restriction in resting macrophages caused by *ribA2* repression can be rescued with exogenous riboflavin, we added 53 μM riboflavin to the cell culture medium. Exogenous riboflavin partially rescued growth of *ribA2* Tet-OFF in THP-1 cells when *ribA2* transcriptional repression was initiated at day 0, but not when *ribA2* was repressed in the pre-culture (**Fig 4C**). These data suggest that *M. tuberculosis* cannot acquire sufficient riboflavin during growth in macrophages, even if it is present at a high level exogenously.

### *M. tuberculosis* requires RibA2 for survival during both acute and chronic phases of infection in mice

To determine the importance of riboflavin biosynthesis during infection, we infected C57BL/6 mice by aerosol with ∼100 CFU of WT *M. tuberculosis* Erdman or *ribA2* Tet-OFF and transcriptionally repressed *ribA2* at different stages of infection (initiation, day 0; acute, day 10; chronic, week 6) by treating mice with doxycycline (dox) (**Fig 5A**). For the WT control, dox treatment with did not significantly alter replication in the lungs, dissemination to the spleen, or persistence (**Fig 5B & 5C**). Without dox treatment, the *ribA2* Tet-OFF strain grew in the lungs, disseminated to the spleen, and persisted in each tissue similarly to the WT control, with only a slight replication defect at 1-2 weeks, during the acute phase of infection (**Fig 5B & 5C**). These data show that the *ribA2* Tet-OFF strain is not highly attenuated without *ribA2* transcriptional repression.

**Figure 5.**
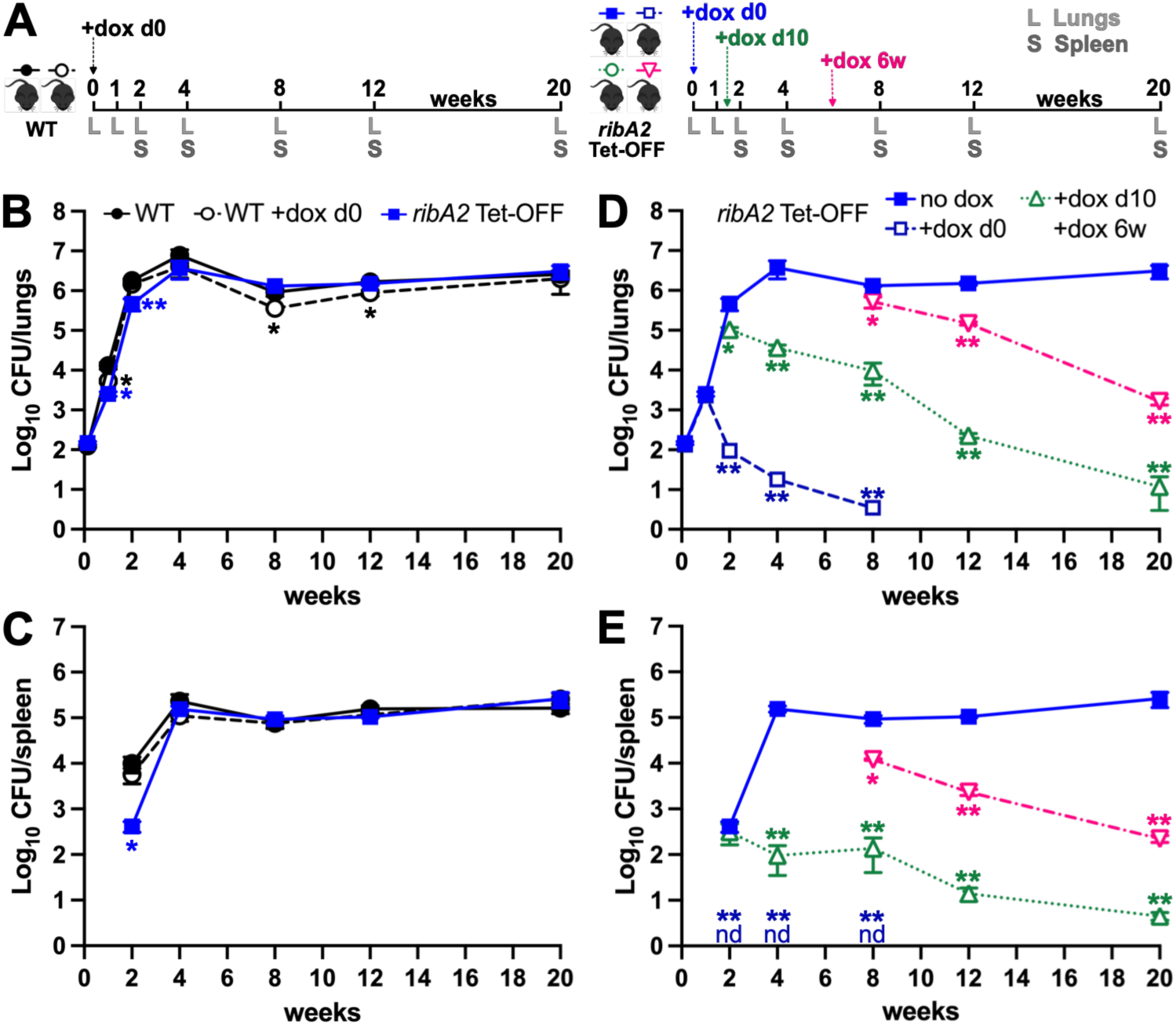
*M. tuberculosis* requires RibA2 for replication and persistence in C57BL/6 mice. **A.** Experiment design. C57BL/6 mice were infected with ∼100 CFU of WT Erdman or *ribA2* Tet-OFF by aerosol. At the indicated times (day 0, day 10 or week 6), groups mice were treated with doxycycline (dox; 2,000 ppm in chow), which was maintained for the duration of the experiment. Groups of mice (*n*=6, 3M, 3F) were euthanized at the indicated time points to collect lungs and spleen. **B-E.** CFU enumeration in lung (**B, D**) and spleen (**C, E**) homogenates by serial dilution and plating on 7H10 agar (WT) or 7H10 with 53 μM riboflavin (*ribA2* Tet-OFF). Data are means ± SEM (*n*=6). Blue “nd” indicates none detected for *ribA2* Tet-OFF +dox d0. Asterisks indicate significant differences from the WT no dox control (**B, C**) or from the *ribA2* Tet-OFF no dox control (**D, E**) by Welch’s *t*-test (24 hr and 1 wk lung data) or by one-way ANOVA with Dunnett’s multiple comparisons test post-hoc. \**P*<0.05, \*\**P*<0.001.

Transcriptional repression of *ribA2* with dox at the start of infection (day 0) did not immediately impact *M. tuberculosis* growth. CFU of the *ribA2* Tet-OFF strain increased in the lungs for the first week in dox-treated mice, like the no dox control (**Fig 5D**). However, after the 1-week time point, lung CFU declined and by 8 weeks post-infection *ribA2* Tet-OFF CFUs were near the detection limit in all mice (**Fig 5D**). We did not detect *ribA2* Tet-OFF bacteria in the spleen at any time point when dox treatment was initiated at day 0 (**Fig 5E**). These data indicate that *M. tuberculosis* requires RibA2 function for sustained replication in the lungs during acute infection. Growth of *ribA2* Tet-OFF during the first week may be due to intracellular flavin stores, as this strain also survives for at least 7 days *in vitro* in medium without riboflavin (**Fig 2B**).

Transcriptional repression of *ribA2* during acute infection (+dox, day 10) caused rapid clearance of *ribA2* Tet-OFF from the lungs and slower clearance from the spleen (**Figs 5D & 5E**). By 20 weeks post-infection, CFU in the lungs and spleens of most mice were at or near the limit of detection (**Fig 5D & 5E**). At the 20 week time point, two of the six dox-treated mice had increased CFU in the lungs and spleen compared to the average CFU at 12 weeks (**Fig S4H & S4K**). Both mice were males, but we did not observe significant differences in CFU between male and female mice in most other experimental groups (**Fig S4**).

We reasoned that the *ribA2* Tet-OFF bacteria in these two mice might have acquired secondary mutations. We picked single colonies isolated from the lungs of each mouse and tested their growth on 7H10 agar without riboflavin +Atc. We identified multiple isolates that grew in this condition (26% of colonies from mouse 1, 83% of colonies from mouse 2) and conducted whole genome sequencing on one such isolate from each mouse. Both mouse isolates harbored a non-synonymous C-A single nucleotide polymorphism in *tetR38* on pGMCS-*ribA2* (**Fig S5A**), which was confirmed by Sanger sequencing (**Fig S5B**). The mutation causes an H44N substitution in the TetR38 DNA binding domain (**Fig S5C**). The *ribA2* Tet-OFF mouse isolates with the *tetR38-H44N* mutation grow in 7H9 +Atc without riboflavin (**Fig S5D**), suggesting that TetR38-H44N cannot repress *ribA2* transcription. We therefore excluded these two mice from the data shown in Fig 5. Our data indicate that *M. tuberculosis* requires RibA2 for growth and survival during acute infection and suggest that there is strong selective pressure to maintain RibA2 expression.

Transcriptional repression of *ribA2* with dox during chronic infection (6 weeks) also led to clearance of *ribA2* Tet-OFF bacteria at a similar rate to that observed during acute infection (**Figs 5D & 5E**). We observed significant reductions in lung and spleen CFU at the 8-week time point (2 weeks of dox treatment) (**Figs 5D & 5E**). By 20 weeks post-infection (14 weeks of dox treatment) CFUs were reduced by approximately 3 logs compared to the no dox control in both lungs and spleens (**Figs 5D & 5E**). These data demonstrate that *M. tuberculosis* requires RibA2 for persistence during chronic infection and suggest that RibA2 inhibitors could function as anti-tubercular agents.

### *M. tuberculosis* requires RibA2 function to survive in mice with necrotic granulomas

We observed that exogenous riboflavin can overcome the requirement for RibA2 *in vitro*, suggesting that *M. tuberculosis* might be able to scavenge sufficient riboflavin from host tissues. The serum concentration of riboflavin (10-33 nM) is several orders of magnitude lower than the 10-20 μM required to rescue growth of riboflavin synthesis deficient *M. tuberculosis* mutants *in vitro* (22, 42). *M. tuberculosis* can replicate extracellularly in necrotic granulomas, a hallmark of active TB disease in humans, which may have a higher riboflavin concentration in the dead cell debris. C57BL/6 mice are a standard model of *M. tuberculosis* infection but fail to form necrotic granulomas (43). C3HeB/FeJ (Kramnik) mice are highly susceptible to *M. tuberculosis* infection and develop necrotic granulomas in which the bacteria can be found extracellularly (44, 45). We therefore used C3HeB/FeJ mice to test whether *M. tuberculosis* requires RibA2 for survival in necrotic granulomas.

We infected C3HeB/FeJ mice with ∼50 CFU of WT *M. tuberculosis* Erdman or the *ribA2* Tet-OFF strain (**Fig 6A**). We initiated *ribA2* transcriptional repression with dox at either 3 weeks post-infection, when organized granuloma lesions begin to develop, or 6 weeks post-infection, when granulomas become necrotic (**Fig 6A**) (45). For the WT control, treatment with dox starting at 3 weeks caused a slight decrease in CFU in the lungs, but increased CFU in the spleen at 6 weeks post-infection (**Figs 6B-C**). Variability in the WT CFU in dox-treated mice was associated with the sex of the animals, with dox-treated female mice tending to have higher CFU, though this was not statistically significant (**Fig S6A-D**).

**Figure 6.**
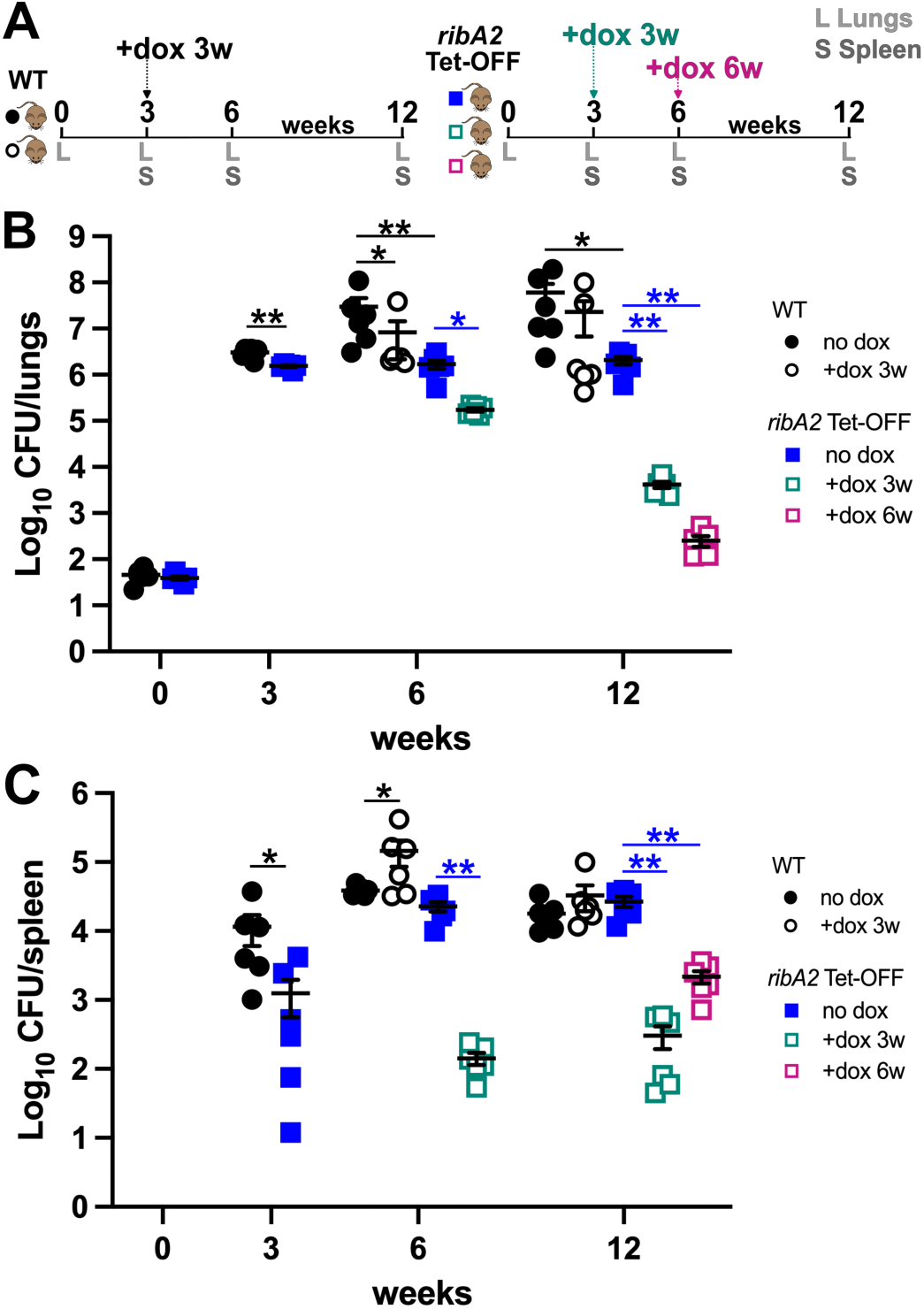
*M. tuberculosis* requires RibA2 for persistence in C3HeB/FeJ mice. **A.** Experiment design. C3HeB/FeJ mice were infected with ∼50 CFU of WT Erdman or *ribA2* Tet-OFF by aerosol. At 3 or 6 weeks post infection, groups of mice were treated with doxycycline (dox, 2,000 ppm in chow) and dox treatment was maintained for the duration of the experiment. Groups of mice (*n*=6, 3M, 3F) were euthanized at the indicated time points to collect lungs and spleen. **B-C.** CFU enumeration in lung (**B**) and spleen (**C**) homogenates by serial dilution and plating on 7H10 agar (WT) or 7H10 with 53 μM riboflavin (*ribA2* Tet-OFF). Data are means ± SEM (*n*=6). Asterisks indicate significant differences compared to the WT (black) or *ribA2* Tet-OFF no dox control (blue) by Welch’s *t*-test (24 hr and 3 wk) or by one-way ANOVA with Dunnett’s multiple comparisons test post-hoc. \**P*<0.05, \*\**P*<0.001.

The *ribA2* Tet-OFF strain had significant defects in growth in the lungs and dissemination to the spleen at 3 weeks post-infection relative to the WT control (**Fig 6B-C**). Lung CFU of the *ribA2* Tet-OFF strain also did not increase further after the 3-week time point, unlike WT *M. tuberculosis* that continued to grow between 3 and 12 weeks post-infection (**Fig 6B**). After delayed dissemination, the *ribA2* Tet-OFF strain achieved similar CFU in the spleen as the WT control at the 6- and 12-week time points (**Fig 6C**). These data suggest that stressors encountered in the C3HeB/FeJ mouse lung restrict replication of the *ribA2* Tet-OFF strain, even without *ribA2* transcriptional repression.

Repression of *ribA2* with dox starting at 3 weeks post-infection led to clearance of *ribA2* Tet-OFF from the lungs and spleen (**Fig 6B-C**). Surprisingly, initiating dox treatment at 6 weeks post-infection led to more rapid clearance of *ribA2* Tet-OFF bacteria from the lungs, with a 4-log reduction in CFU by 12 weeks (**Fig 6B**). Dox treatment starting at 6 weeks post infection also caused a nearly 2-log reduction in spleen CFU by 12 weeks (**Fig 6C**). These data suggest that loss of RibA2 function increases *M. tuberculosis* susceptibility to stressors encountered in the necrotic lung lesions of C3HeB/FeJ mice and that *M. tuberculosis* does not have access to sufficient riboflavin to sustain its viability within these lesions.

## DISCUSSION

New targets for anti-tubercular drug development must be identified to combat emerging drug resistance and reduce TB treatment duration. Here, we show that *M. tuberculosis* requires the riboflavin biosynthesis enzyme RibA2 for survival *in vitro* in the absence of exogenous riboflavin, for growth in resting and IFN-γ activated macrophages and for survival at all stages of infection in two mouse infection models. These include C3HeB/FeJ mice that develop necrotic lung granulomas, which mimic active TB disease in humans. Further, we show that inhibition of riboflavin biosynthesis by repression of *ribA2* triggers changes to the *M. tuberculosis* transcriptome that indicate decreased electron transport chain function and broad changes to central carbon and lipid metabolism. These transcriptional adaptations are consistent with reduced function of known flavin-dependent enzymes. Our data are important because they establish RibA2 as a highly vulnerable target for development of new anti-tubercular drugs in relevant animal models of active TB disease.

Transcriptional repression of *ribA2* was bactericidal to *M. tuberculosis* in the absence of exogenous riboflavin. Our data are consistent with a genome-wide CRISPRi screen that identified *ribA2* as highly vulnerable to transcriptional inhibition, like the targets of existing anti-tubercular drugs (14). Our data are also consistent with the observation that CRISPRi-mediated transcriptional silencing of *ribA2*, *ribG*, *ribH* or *ribF* caused death of *M. tuberculosis* in media without riboflavin (22). We observed slower death of our *ribA2* Tet-OFF strain over the first week of culture in the absence of riboflavin as compared to the study that used CRISPRi to silence *ribA2* (22). This may be due to differences in the method for recovering surviving bacteria. The previous study used standard Middlebrook 7H10 agar without riboflavin (22), but we recovered the *ribA2* Tet-OFF strain on agar containing riboflavin, to enable regrowth of bacteria that were depleted of flavins. Our data demonstrate that RibA2 is essential for *M. tuberculosis* growth and survival in standard culture conditions.

Our *ribA2*::Tn and *ribC*::Tn mutants exhibited a slow rate of death in the absence of riboflavin compared to *ribA2* Tet-OFF cultured with Atc to repress *ribA2*. Slow death in the absence of riboflavin was also observed for a Δ*ribC* mutant (46). The Tn mutants and Δ*ribC* must be grown with riboflavin. *M. tuberculosis* grown with riboflavin have a 50- to 100-fold higher intracellular riboflavin concentration and exhibit increased abundance of the flavin-sequestering proteins Acg and dodecin (47–51). Intracellular riboflavin stores could promote survival of the *ribA2*::Tn and *ribC*::Tn mutants when they are shifted to medium without riboflavin. Since we grew *ribA2* Tet-OFF without riboflavin prior to repression of *ribA2* with Atc, it is expected to lack these flavin stores. However, FMN and FAD are highly stable in bacteria (52). FMN and FAD bind tightly to flavoproteins and do not dissociate with each catalytic cycle (19), which may limit their turnover. Flavin stability could contribute to survival of the *ribA2* Tet-OFF strain during the first week after *ribA2* repression both *in vitro* without riboflavin and in mice. Thus, intracellular riboflavin stores may temporarily sustain viability of *M. tuberculosis* that cannot synthesize riboflavin.

Riboflavin starvation ultimately kills *M. tuberculosis*, and our transcriptional profiling data provide insight into the molecular mechanisms underlying this loss of viability. We observed increased expression of genes encoding the Cyt-*bd* and NDH-2 (*ndh*) electron transport chain components when *ribA2* was repressed. Increased expression of Cyt-*bd* and *ndh* was also observed in *M. tuberculosis* mutants lacking cytochrome-*bc_1_*-*aa_3_* oxidase (53) and in response to cytochrome *c* oxidase inhibitors (54), both of which decrease electron transport chain activity. Depletion of FMN and FAD is predicted to decrease activity of the NADH dehydrogenases NHD-1 (*nuoA-N*) and NDH-2 (*ndh* and *ndhA*), leading to lower electron transport chain activity. Loss of both *ndh* and *ndhA* is synthetically lethal in standard culture conditions *in vitro*, so decreased NDH-2 activity alone could cause death of *M. tuberculosis* (41, 53). A *M. tuberculosis* mutant lacking NHD-2 activity was killed by chemical inhibitors of NDH-1 due to an inability to oxidize NADH to NAD^+^ (53, 55). Chemical inhibition of both NDH-1 and NDH-2 also killed *M. tuberculosis* (55). Thus, indirect inhibition of NADH dehydrogenases by depletion of essential flavin cofactors could cause death of *M. tuberculosis* when riboflavin biosynthesis is inhibited.

Other metabolic effects of flavin cofactor depletion could also kill *M. tuberculosis* when riboflavin biosynthesis is inhibited. We observed increased expression of many *fadE* genes when *ribA2* was transcriptionally repressed. FadE enzymes are FAD-dependent acyl-CoA dehydrogenases (ACADs) that catalyze a step in fatty acid or cholesterol β-oxidation (56). Reoxidation of the FAD cofactor in ACADs depends on a short electron transport pathway comprised of EtfD and EtfBA (35). *M. tuberculosis* requires EtfD for growth in fatty acid containing media *in vitro* and for growth and survival in aerosol-infected mice (35). Thus, loss of ACAD function due to decreased FAD cofactor availability could also cause death of *M. tuberculosis* that cannot synthesize riboflavin. This may be particularly relevant in the infected host, in which *M. tuberculosis* uses fatty acids and cholesterol as carbon sources (31, 57).

Transcriptional repression of *ribA2* prevented *M. tuberculosis* growth in both resting and IFN-γ activated monocyte-derived macrophages, and this growth inhibition could not be overcome by exogenous riboflavin. Our data suggest that *M. tuberculosis* cannot acquire sufficient riboflavin from macrophages and must instead synthesize riboflavin to supports its intracellular replication. The riboflavin concentration in multiple cultured mammalian cell types is 1-10 μM (58), which is lower than the ∼20 μM riboflavin needed to rescue growth of riboflavin synthesis deficient *M. tuberculosis in vitro* (22). We were surprised that IFN-γ activated macrophages did not efficiently kill *M. tuberculosis* when *ribA2* was repressed, but human monocyte-derived macrophages have a relatively low response to IFN-γ stimulation (59). It will be interesting to determine how riboflavin synthesis inhibition impacts survival of *M. tuberculosis* in other infected cell types, such as alveolar macrophages and neutrophils. Our experiments also revealed that *M. tuberculosis* requires RibA2 for survival in mice at all stages of infection, suggesting low bioavailability of riboflavin in infected host tissues. This is consistent with the low concentration of riboflavin (10-33 nM) in human serum (42). Our data support that *M. tuberculosis* must synthesize riboflavin to enable its survival during infection.

We observed clearance of *ribA2* Tet-OFF after *ribA2* transcriptional repression in both C57BL/6 mice that form cellular lung lesions and C3HeB/FeJ mice that form hypoxic necrotic lung granulomas. To our knowledge, our study is the first to use tetracycline regulated conditional expression to demonstrate essentiality of a *M. tuberculosis* gene in the C3HeB/FeJ model. These mice are a useful model because they have an elevated type I interferon response and develop necrotic lung granulomas like humans with active TB. We observed modest growth inhibition of WT *M. tuberculosis* Erdman in the lungs of C3HeB/FeJ mice treated with dox and high mouse-to-mouse variability. Previous studies have shown that dox accumulates in necrotic lesions in rabbits and mice (60, 61), suggesting that variability is not due to low dox tissue penetration. Instead, differences in the response to dox may be due to variability in lung granuloma lesion development, which influences both mouse survival and antibiotic treatment efficacy (45, 62, 63). Nevertheless, our data indicate that inhibition of riboflavin biosynthesis can effectively clear *M. tuberculosis* from necrotic granulomas, which are associated with active TB disease in humans.

We observed more rapid clearance of our *ribA2* Tet-OFF strain from the lungs of C3HeB/FeJ when we initiated treatment with dox at a late time point, after necrotic granulomas develop. *M. tuberculosis* that cannot synthesize riboflavin may be particularly sensitive to host stress in these lesions. Granulomas in C3HeB/FeJ mice are enriched in neutrophils that produce neutrophil extracellular traps (64–66) and are hypoxic (67). Riboflavin biosynthesis inhibition is expected to sensitize *M. tuberculosis* to oxidative stress, since many redox active enzymes depend on flavin or deazaflavin cofactors (16, 21). The hypoxic environment in granulomas is also expected to reduce flavin cofactors, which would broadly impact metabolism by inactivating many flavin-dependent enzymes (68). *M. tuberculosis* in hypoxic granulomas are generally more tolerant to antibiotics (61, 69, 70), but our data suggest they would be sensitized to riboflavin biosynthesis inhibitors. Further studies will be needed to identify the host immune mechanisms that mediate clearance of *M. tuberculosis* when riboflavin biosynthesis is inhibited.

Although inhibition of riboflavin biosynthesis enables the host to clear *M. tuberculosis* infection, it may also reduce recognition of *M. tuberculosis* infected cells by mucosal invariant T (MAIT) cells. Riboflavin biosynthesis intermediates are key activating ligands of MR1-restricted MAIT cells (71, 72). MAIT cells are enriched in the airways of humans with TB and contribute to the early control of *M. tuberculosis* infection in mice (73, 74). Several human MAIT cell clones showed reduced recognition of a *M. tuberculosis* Δ*ribA2* mutant, consistent with reduced production of MAIT cell activating riboflavin metabolites (47). Transcriptional repression of *ribA2* is also expected to reduce MAIT cell recognition of *M. tuberculosis* infected cells. Since we observed clearance of *ribA2* Tet-OFF from mice after *ribA2* repression, our data suggest that MAIT cells specific for riboflavin metabolites play a limited role in killing *M. tuberculosis* when riboflavin biosynthesis is inhibited. However, it may be possible to further improve *M. tuberculosis* control by inhibiting steps in riboflavin biosynthesis that cause accumulation of MAIT cell activating intermediates. We will test this idea in the future using *M. tuberculosis* strains that conditionally express other riboflavin biosynthesis enzymes.

Our data support the riboflavin biosynthesis pathway as a promising target for development of new antibiotics to treat TB. Inhibition of RibA2 is predicted to limit production of all riboflavin biosynthesis intermediates, including those detected by MR1-restricted MAIT cells, which may reduce MAIT cell-mediated control of *M. tuberculosis* infection. Inhibition of RibA2 is also predicted to prevent production of the F_420_ deazaflavin, a cofactor for the Ddn nitroreductase that activates the anti-tubercular pro-drugs pretomanid and delamanid (75, 76). The riboflavin biosynthesis enzymes RibC and RibH that are not required to produce MAIT cell agonists or F_420_ may offer greater promise as new antibiotic development targets since inhibitors of these enzymes would not interfere with MAIT cell recognition or the activity of other antibiotics. Our future studies will include use of conditional expression technologies to determine whether these other riboflavin biosynthesis enzymes are equally vulnerable to inhibition during infection as RibA2. Importantly, the riboflavin biosynthesis pathway is highly conserved in other pathogenic mycobacteria, so inhibitors targeting this pathway could function as broad-spectrum antibiotics to treat TB and other mycobacterial infections.

## MATERIALS AND METHODS

### Bacterial strains and culture conditions

Bacterial strains are listed in **Table S2**. *M. tuberculosis* Erdman and the *ribA2* Tet-OFF derivative strain were grown at 37°C in Middlebrook 7H9 (Difco) liquid medium supplemented with 10% albumin-dextrose-saline (ADS), 0.5% glycerol and 0.1% Tween-80 with aeration or on Middlebrook 7H10 agar (Difco) supplemented with 10% oleic acid-albumin-dextrose-catalase (OADC; BD Biosciences) and 0.5% glycerol unless otherwise noted. *M. tuberculosis* Erdman *ribA2*::Tn and *ribC*::Tn mutants isolated from our arrayed Tn mutant library (27) were grown in media that was additionally supplemented with 53 μM riboflavin. Frozen stocks were made by adding glycerol at 15% final concentration to cultures grown to late exponential phase and were stored at -80°C. Antimicrobials were used at the following concentrations: streptomycin (Str) 25 μg/ml, hygromycin (Hyg) 50 μg/ml, kanamycin (Kan) 15 μg/ml, and cycloheximide 100 μg/ml.

### Cloning and strain construction

The *ribA2* Tet-OFF plasmid pGMCS-*ribA2* was constructed using the previously described Gateway cloning strategy (77). Briefly, *M. tuberculosis* Erdman *ribA2* with its native ribosome binding site was amplified by PCR using the primers listed in **Table S3** and inserted in the pDO23A entry vector with BP clonase (Invitrogen) to create pEN23A-*ribA2*. The pGMCS-*ribA2* construct was assembled by using LR clonase (Invitrogen) with pEN23A-*ribA2*, pEN12A-P750, pEN41A-T38S38 and pDE43-MCS. The pGMCS-*ribA2* construct was confirmed by whole plasmid sequencing (Plasmidsaurus, Louisville, KY). WT *M. tuberculosis* Erdman was electroporated with pGMCS-*ribA2* as described (78) and transformants were selected on 7H10 agar containing Str. Integration of the pGMCS-*ribA2* plasmid was confirmed by PCR with primers listed in **Table S3**.

The pJG1100-*ΔribA2* vector was created by PCR amplifying ∼900 bp regions of the *M. tuberculosis* Erdman chromosome flanking *ribA2* using primers listed in **Table S3**, digesting the resulting products with *Pac*I/*Avr*II or *Avr*II/*Asc*I and ligating these fragments in the pJG1100 suicide vector (79) digested with *Pac*I/*Asc*I. The pJG1100-Δ*ribA2* construct was confirmed by whole plasmid sequencing (Plasmidsaurus, Louisville, KY). The pJG1100-Δ*ribA2* plasmid was electroporated into *M. tuberculosis* carrying pGMCS-*ribA2* and transformants were selected on 7H10 agar containing 53 μM riboflavin, Str, Kan and Hyg. Transformants with the pJG1100-Δ*ribA2* plasmid integrated were confirmed by PCR using primers listed in **Table S3**, then grown in 7H9 broth with 53 μM riboflavin and Str to mid-exponential phase to allow excision of pJG1100-Δ*ribA2* and plated on 7H10 agar containing 53 μM riboflavin, Str and 2% sucrose for counter-selection. Colonies were grown from 7H10 sucrose plates in 7H9 broth with 53 μM riboflavin and Str and screened for the Δ*ribA2* deletion by PCR (**Table S3**).

### Phthiocerol dimycocerosate (PDIM) assay

*M. tuberculosis* cultures grown in 10 ml complete 7H9 medium to mid-logarithmic phase were labeled for 48 hr with 10 μCi of [1-^14^C] propionic acid, sodium salt (American Radiochemicals, Inc; specific activity 50-60 mCi/mmol) and apolar lipids were extracted as described (79). Labeled lipids were separated by thin layer chromatography and detected by phosphor imaging as described (27).

### Whole genome sequencing

Genomic DNA was extracted by the CTAB-lysozyme method from *M. tuberculosis ribA2* Tet-OFF and two *ribA2* Tet-OFF mouse isolates as described (80). DNA was cleaned with the Genomic DNA clean and concentrator kit 25 (Zymo) and submitted to SeqCoast Genomics (*ribA2* Tet-OFF; Portsmouth, NH) or SeqCenter (*ribA2* Tet-OFF mouse isolates; Pittsburgh, PA) for library preparation using the Illumina DNA Prep tagmentation kit with unique dual indices and sequencing on the Illumina NextSeq 2000 using a 300 cycle flow cell kit (2×150 bp reads) with 1-2% PhiX control spiked in to optimize base calling. Read demultiplexing, trimming and run analytics were performed using DRAGEN v3.10.12. To generate consensus sequences for *ribA2* Tet-OFF and each mouse isolate, paired reads were mapped to the *M. tuberculosis* Erdman reference genome (NC_020559.1) using the “map to reference” function in Geneious Prime 2022 software (Biomatters, Ltd.) as described (27). The *ribA2* Tet-OFF genome was examined for single nucleotide polymorphisms (SNPs) in the locus required for PDIM production (*tesA-lppX*). The *ribA2* Tet-OFF mouse isolate consensus genomes were aligned to the *ribA2* Tet-OFF parental consensus to identify SNPs using Mauve alignment in Geneious Prime with the default settings. To identify SNPs in pGMCS-*ribA2* in the mouse isolates, sequencing reads were mapped to pGMCS-*ribA2* using the “map to reference” function and the resulting consensus sequences were aligned to pGMCS-*ribA2* using Mauve alignment in Geneious Prime.

### Growth curves

Bacteria were grown to mid-exponential phase (OD_600_ = 0.5) in complete 7H9 medium (WT, *ribA2* Tet-OFF) or 7H9 with 53 μM riboflavin (*ribA2*::Tn, *ribC*::Tn), then inoculated in triplicate in complete 7H9 medium ± 53 μM riboflavin and ± 100 ng/ml anhydrotetracycline (Atc). Fresh Atc was added to cultures every 7 days to maintain *ribA2* repression in *ribA2* Tet-OFF. The OD_600_ of cultures was measured every 2-3 days. Cultures were serially diluted in phosphate buffered saline (PBS) containing 0.05% Tween-80 (PBS-T) and plated on 7H10 (WT) or 7H10 with 53 μM riboflavin (*ribA2* Tet-OFF, *ribA2*::Tn, *ribC*::Tn) every 3-5 days. Plates were incubated at 37°C for at least 4 weeks before counting CFUs.

### RNAseq transcriptional profiling

WT and *ribA2* Tet-OFF were grown in complete 7H9 to mid-logarithmic phase (OD_600_ ∼0.5), then diluted to OD_600_ = 0.2 in 10 ml 7H9 ± 100 ng/ml Atc or 7H9 + 0.1% ethanol (solvent control). Triplicate cultures prepared for each time point and condition were collected for RNA extraction on days 1, 2 and 4 (*ribA2* Tet-OFF) or day 1 (WT). Bacteria were pelleted by centrifugation (4,100 x*g*, 10 min, 4°C) and RNA was extracted in Trizol (Ambion) by bead beating as described (81). RNA purity was assessed using a Nanodrop 2000 (ThermoScientific). RNA was submitted to the University of Minnesota Genomics Center for quantification with RiboGreen and Agilent TapeStation sizing, library preparation with the Illumina Stranded Total RNA with Ribozero plus kit, and sequencing on the Element AVITI Freestyle (2×150 bp paired end).

Sequence reads were analyzed using CHURP bulk_rnaseq (82), developed and maintained by the Minnesota Supercomputing Institute. For each paired-end read, the CHURP pipeline is as follows: summarize read quality with fastqc, filter low-quality reads and trim adapters using trimmomatic, map reads to the genome using hisat2 to create BAM files, indexing and sorting with samtools, and using featurecounts to create a counts matrix for DESeq2 analyses. A hisat2 index for read mapping was created for the Erdman genome using the GCF_049959555.1.fasta file. Sequence reads corresponding to rRNA and tRNA were overrepresented based on fastqc analysis, so a modified Erdman genome GCF_049959555.1.gtf annotated file was created with rRNA and tRNA sequences removed and was subsequently used with featurecounts to annotate mapped reads, exclude reads that mapped to rRNA and tRNA, and create matrix counts tables. Similar analyses were done using the CHURP bulk_rnaseq pipeline to create matrix counts tables against the H37Rv genome to enable annotation of Rv gene numbers. For these analyses, a hisat2 index for H37Rv GCF_026185275.1.fasta was created along with a modified H37Rv genome annotation file GCF_26185275.1.gtf with rRNA and tRNA sequences removed for featurecounts.

The DESeq2 (83) Bioconductor package in R (84) was used to determine differential expression for all genes in the matrix counts files for both Erdman and H37Rv mappings. PCA conducted by DESeq2 confirmed high reproducibility of biological replicates for both genome mapping strategies. Volcano plots were constructed in R for both genome mappings with similar results. Heat maps were made in GraphPad Prism (v11.1.0 for Mac; GraphPad Software, Boston, MA) using differential expression data produced by DESeq2. R was used to analyze DESeq2 differential gene expression data to identify differentially expressed flavoprotein genes based on a curated list of *M. tuberculosis* H37Rv flavoproteins from UniProt (UP000001584) using an adjusted *P* value (*Padj*) <0.05 and log_2_ fold change > ±1. Flavoproteins were assigned to a flavin class (e.g. FAD-binding, FMN-binding, flavin biosynthesis or indirect) based on UniProt classifications, associated publications and protein homology. Significantly up- or down-regulated genes were assigned to a flavin class and ranked by expression level within the flavin class. Divergent flavin class distributions and ranked gene expression levels showing log_2_ fold change were plotted using ggplot in R.

### Quantitative RT-PCR

RNA extracted for RNAseq was treated with Turbo DNase (Invitrogen) and converted to cDNA with the Roche Transcriptor First Strand kit (Roche) using random hexamers for priming as described (81). No RT controls were included for each sample to assess genomic DNA contamination. The *ribA2*, *ribH*, *cydA*, *frdA* and *16S rRNA* transcripts were detected by quantitative PCR using 2.5 μM of each primer (**Table S3**) and the 2x SYBR green master mix (Roche) on a LightCycler 480 (Roche) using technical duplicates for each sample as described (81). Cycle threshold values (Cp) were converted to copy numbers using standard curves generated using genomic DNA for each primer set. All no RT controls indicated no appreciable genomic DNA contamination. Transcript copy numbers were normalized to the *16S rRNA* copy number for the same sample.

### Growth in THP-1 cells

The THP-1 (ATCC TIB-202) human monocytic cell line was cultured in RPMI base (Gibco) supplemented with non-heat treated 10% fetal bovine serum (Corning), hereafter referred to as RPMI, in a humidified incubator at 37°C with 5% CO_2_. THP-1 cells were differentiated into monocyte-derived macrophages with 50 nM phorbol 12-myristate-13-acetate (PMA, Sigma) for 24 hr in 96-well tissue culture plates. Differentiated cells were washed twice with Hanks Balance Salt Solution (Corning), given fresh RPMI and rested for 72 hr before infection. Rested macrophages were activated with IFN-γ using RPMI containing 20 ng/ml human IFN-γ (Peprotech) for an additional 24 hr and then infected.

Bacteria were grown in complete 7H9 from frozen stocks to mid-exponential phase (OD_600_ = 0.4-0.7). For preculture with Atc, strains were grown for three days with 100 ng/ml Atc prior to starting the infection. Bacteria were collected by centrifugation (2,500 x*g*, 10 min), resuspended in PBS containing 0.05% Tween-80 (PBS-T), de-clumped by centrifugation (50 x*g*, 5 min) and diluted in RPMI at the appropriate density. Infections were initiated at a 1:20 bacteria:macrophage multiplicity of infection (MOI) in fresh RPMI ± 200 ng/ml Atc for 2 hr at 37°C with 5% CO_2_. Extracellular bacteria were removed by three washes with pre-warmed Dulbecco’s PBS (Gibco). Fresh pre-warmed RPMI ± 200 ng/ml Atc or RPMI + 53 μM riboflavin ± 200 ng/ml Atc was added and cells were incubated at 37°C with 5% CO_2_. The cell culture medium was replaced every 2-3 days. Viable intracellular bacteria were enumerated at 2 hr, 1, 3 and 7 days post-infection. THP-1 cells were lysed with 0.1% Tween-80 at 37°C for 10 min. Monolayers were visually inspected for lysis before serially diluting the lysate in PBS-T and plating on 7H10 (WT) or 7H10 with 53 μM riboflavin (*ribA2* Tet-OFF). Plates were incubated for at least 4 weeks at 37°C before counting CFU.

### Mouse infections

Male and female C57BL/6J or C3HeB/FeJ mice 6 weeks of age were purchased from The Jackson Laboratory, USA. Mice were infected by the aerosol route with ∼100 CFU (C57BL/6J) or ∼50 CFU (C3HeB/FeJ) of WT *M. tuberculosis* Erdman or *ribA2* Tet-OFF using an Inhalation Exposure System (GlasCol) as described (85). Bacteria were grown to mid-exponential phase (OD_600_ of ∼0.5) in complete 7H9, collected by centrifugation (2,850 x*g*, 10 min), washed once in PBS-T, de-clumped by centrifugation (58 x*g*, 5 min), and diluted to an appropriate OD_600_ in PBS-T to achieve the desired input dose: WT OD_600_ = 0.007 for 100 CFU, OD_600_ = 0.002 for 50 CFU; *ribA2* Tet-OFF OD_600_ = 0.012 for 100 CFU, OD_600_ = 0.003 for 50 CFU. Groups of mice were treated with doxycycline (dox) chow (Teklad Global 2018 diet containing 2,000 ppm dox; Envigo) starting at the following times: day 0 (immediately after infection), day 10 or day 42 (week 6) for C57BL/6 mice; day 21 (week 3) or day 42 (week 6) for C3HeB/FeJ mice. Dox chow was stored at 4°C until use and was replaced weekly. Groups of mice (*n*=6, 3 male, 3 female) were euthanized by CO_2_ overdose at the following time points: 24 hr, 1, 2, 4, 8, 12 and 20 weeks post-infection for C57BL/6 mice; 24 hr, 3, 6, and 12 weeks post-infection for C3HeB/FeJ mice. Lungs were recovered at 24 hr and 1 week; at all other time points both lungs and spleens were collected. Organs were homogenized in PBS-T, serially diluted in PBS-T and plated on 7H10 (WT) or 7H10 with 53 μM riboflavin (*ribA2* Tet-OFF) containing cycloheximide. Colonies were counted after at least 5 weeks of incubation at 37°C.

### Animal ethics statement

All animal protocols used in this study were reviewed and approved by the Institutional Animal Care and Use Committee of the University of Minnesota under protocol number 2210-40465A. The University of Minnesota’s NIH Animal Welfare Assurance Number is D16-00288 (A3456-01), expiration date 4/30/2028. All animal experiments were conducted in accordance with the recommendations in the Guide for the Care and Use of Laboratory Animals of the National Institutes of Health (86).

### Statistical analyses

For all growth curve data and CFU data from macrophages or mice, data were log_10_ transformed prior to statistical analyses. One-way ANOVA with Dunnett’s multiple comparisons test applied post-hoc was used for comparisons of multiple strains. Comparisons between two groups were done by unpaired *t*-test (+Atc vs no Atc control; males vs females) or by Welch’s *t*-test (WT vs *ribA2* Tet-OFF no dox mouse data) to account for unequal variances (87). *P* values were calculated in GraphPad Prism (v11.1.0 for Mac, GraphPad Software, Boston, MA). *P* values <0.05 were considered significant.

## Data availability

Raw sequencing data from whole genome sequencing and RNAseq experiments are available in FASTA format from the NCBI SRA under BioProject accession numbers PRJNA1534021 and PRJNA1534559, respectively.

## ACKNOWLEDGEMENTS

This work was supported by NIH grants R21AI146440 (ADT), R21AI187844 (ADT) and a University of Minnesota Doctoral Dissertation Fellowship (SNB). Portions of this work were completed with resources, including instrumentation and staff, at the University of Minnesota Genomics Center (RRID: SCR_012413), the University of Minnesota Imaging Centers (RRID: SCR_020997) and the Minnesota Supercomputing Institute (MSI) at the University of Minnesota. The authors thank Dirk Schnappinger for providing the pDO23A, pEN12A-P750, pEN41A-T38S38 and pDE43-MCS plasmids that were used to construct pGMCS-*ribA2*.

## SUPPLEMENTAL FIGURE LEGENDS

**Figure S1.** Construction and validation of *M. tuberculosis ribA2* Tet-OFF. **A.** Diagram of the *ribA2* Tet-OFF genome (Δ*ribA2* pGMCS-*ribA2*) indicating the in-frame, unmarked deletion of *ribA2* and integration of pGMCS-*ribA2* encoding tetracycline-repressible *ribA2* at the phage L5 *attB* site. Small arrows indicate primers. **B.** PCR confirmation of the Δ*ribA2* deletion and integration of pGMCS-*ribA2* in *ribA2* Tet-OFF. **C.** Thin layer chromatograph of ^14^C-labelled apolar lipids extracted from WT and *ribA2* Tet-OFF strains. The DIM A and DIM B forms of the PDIM lipid are indicated. **D-E.** Quantitative RT-PCR. WT Erdman and *ribA2* Tet-OFF were grown in 7H9 medium, diluted to OD_600_ = 0.2 and then grown for 1 or 4 days in 7H9 ± 100 ng/ml anhydrotetracycline (Atc) prior to RNA extraction. Transcript abundance of *ribA2* (**D**) and *ribH* (**E**) were determined relative to 16S rRNA by qRT-PCR. Data are means ± standard deviations of triplicate cultures. Black asterisks indicate statistically significant differences compared to WT no Atc by one-way ANOVA; colored asterisks indicate statistically significant differences for +Atc compared to the corresponding no Atc control by unpaired *t*-test: \**P*<0.05 \*\**P*<0.001.

**Figure S2.** Differential gene expression in WT or *ribA2* Tet-OFF *M. tuberculosis* at days 1 and 2. RNA was extracted from WT Erdman or *ribA2* Tet-OFF grown in 7H9 ± 100 ng/ml Atc or in 7H9 +0.1% ethanol (solvent control) and transcriptional profiles were determined by RNAseq. Volcano plots of log_2_-fold change and adjusted *P*-value determined by RNAseq. Each dot represents an individual gene. Dashed lines indicate the log_2_-fold change = ± 1 and *Padj* < 0.05 statistical significance cutoffs. Red or blue dots represent significantly up- or down-regulated genes relative to the control, respectively. **A.** WT +Atc vs WT no Atc control, day 1. **B.** WT +ethanol vs WT no Atc control, day 1. **C.** *ribA2* Tet-OFF +Atc vs *ribA2* Tet-OFF no Atc, day 1. **D.** *ribA2* Tet-OFF +Atc vs *ribA2* Tet-OFF no Atc, day 2.

**Figure S3.** Lipid metabolism, ribosomal protein genes and flavin-associated genes that are differentially expressed upon *ribA2* transcriptional repression. **A-B.** Heat maps showing the log_2_-fold change in transcript abundance of differentially expressed genes related to fatty acid utilization, lipid synthesis and ribosomal proteins relative to the WT no Atc control (**A**) or relative to the *ribA2* Tet-OFF no Atc control from the same time point (**B**). **C.** Genes encoding flavoproteins that were significantly differentially expressed in *ribA2* Tet-OFF +Atc day 4 relative to WT no Atc control.

**Figure S4.** *M. tuberculosis* lung and spleen burdens from individual C57BL/6 mice, separated by sex. Data from C57BL/6 mice that were infected by aerosol with WT Erdman (**A, B, D, E**) or *ribA2* Tet-OFF (**C, F, G-L**) and treated with doxycycline (dox, 2,000 ppm in chow) starting at the indicated time points, from Fig 5. *M. tuberculosis* CFU in lungs (**A-C, G-I**) and spleen (**D-F, J-L**) were determined by plating. Data from male and female mice (*n*=3 each) are shown separately. Lines indicate the mean. In **J**, nd indicates no CFU detected in any mouse (detection limit = 3 CFU). Asterisks indicate a statistically significant difference in CFU between male and female mice by unpaired *t*-test, \**P*<0.05. In **H** and **K**, X’s over the data for two males indicate mice that were excluded from the averaged data in Fig 5 because *ribA2* Tet-OFF isolates recovered from these mice harbored mutations in *tetR38* (see **Figure S5**).

**Figure S5.** Mutations in *tetR38* prevent conditional repression of *ribA2* in *ribA2* Tet-OFF isolates from two C57BL/6 mice. **A.** Whole genome sequencing reads for the *ribA2* Tet-OFF parental strain and two mouse isolates indicating a consensus C to A single nucleotide substitution in codon 44 of the *tetR38* on pGMCS-*ribA2* in both mouse isolates. **B.** Sanger sequencing confirmation of the A substitution in codon 44 of *tetR38* in both mouse isolates. **C.** Ribbon diagrams of TetR38 (left) and TetR38-H44N (right) protein structures, with the amino acid substitution in the DNA binding domain highlighted. **D.** Growth in complete 7H9 without riboflavin. The *ribA2* Tet-OFF parent strain and *ribA2* Tet-OFF mouse isolates (M1, M2) were grown in complete 7H9 to mid-log phase, then diluted to OD_600_ = 0.01 in 7H9 ± 100 ng/ml anhydrotetracycline (Atc). Growth was monitored by OD_600_ measurement. Data are means ± standard deviations of triplicate cultures. Asterisks indicate statistically significant differences compared to the *ribA2* Tet-OFF parental no Atc control by one-way ANOVA with Dunnett’s multiple comparisons test: \**P*<0.05.

**Figure S6.** *M. tuberculosis* lung and spleen burdens from C3HeB/FeJ mice, separated by sex. Data from C3HeB/FeJ mice that were infected by aerosol with WT Erdman (**A-D**) or *ribA2* Tet-OFF (**E-J**) and treated with doxycycline (dox, 2,000 ppm in chow) starting at the indicated time points, from Fig 6. *M. tuberculosis* CFU in lungs (**A, B, E-G**) and spleen (**C, D, H-J**) were determined by plating. Data from male and female mice (*n*=3 each) are shown separately. Lines indicate the mean. No statistically significant differences between male and female mice were identified by unpaired *t*-test.

## Notes

### Competing Interest Statement

The authors have declared no competing interest.

