## Supplemental Figures S1-S6 for "*Mycobacterium tuberculosis* requires riboflavin biosynthesis for survival in the host"

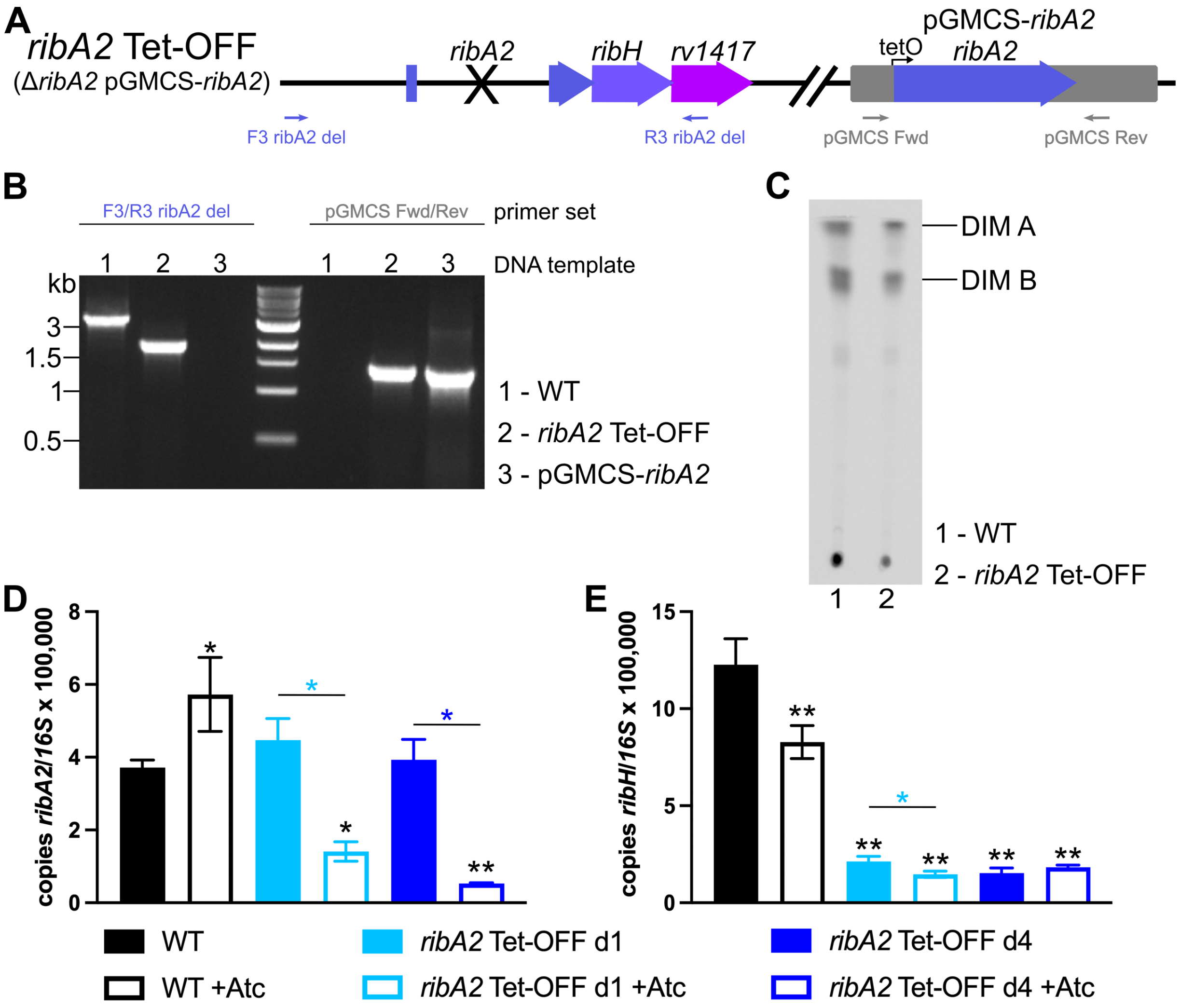


**Figure S1. Construction and validation of *M. tuberculosis ribA2* Tet-OFF. A.** Diagram of the *ribA2* Tet-OFF genome (Δ*ribA2* pGMCS-*ribA2*) indicating the in-frame, unmarked deletion of *ribA2* and integration of pGMCS-*ribA2* encoding tetracycline-repressible *ribA2* at the phage L5 *attB* site. Small arrows indicate primers. **B.** PCR confirmation of the Δ*ribA2* deletion and integration of pGMCS-*ribA2* in *ribA2* Tet-OFF. **C.** Thin layer chromatograph of $^{14}$C-labelled apolar lipids extracted from WT and *ribA2* Tet-OFF strains. The DIM A and DIM B forms of the PDIM lipid are indicated. **D-E.** Quantitative RT-PCR. WT Erdman and *ribA2* Tet-OFF were grown in 7H9 medium, diluted to $OD_{600}$ = 0.2 and then grown for 1 or 4 days in 7H9 ± 100 ng/ml anhydrotetracycline (Atc) prior to RNA extraction. Transcript abundance of *ribA2* (**D**) and *ribH* (**E**) were determined relative to 16S rRNA by qRT-PCR. Data are means ± standard deviations of triplicate cultures. Black asterisks indicate statistically significant differences compared to WT no Atc by one-way ANOVA; colored asterisks indicate statistically significant differences for +Atc compared to the corresponding no Atc control by unpaired *t*-test: *$P$<0.05 **$P$<0.001.

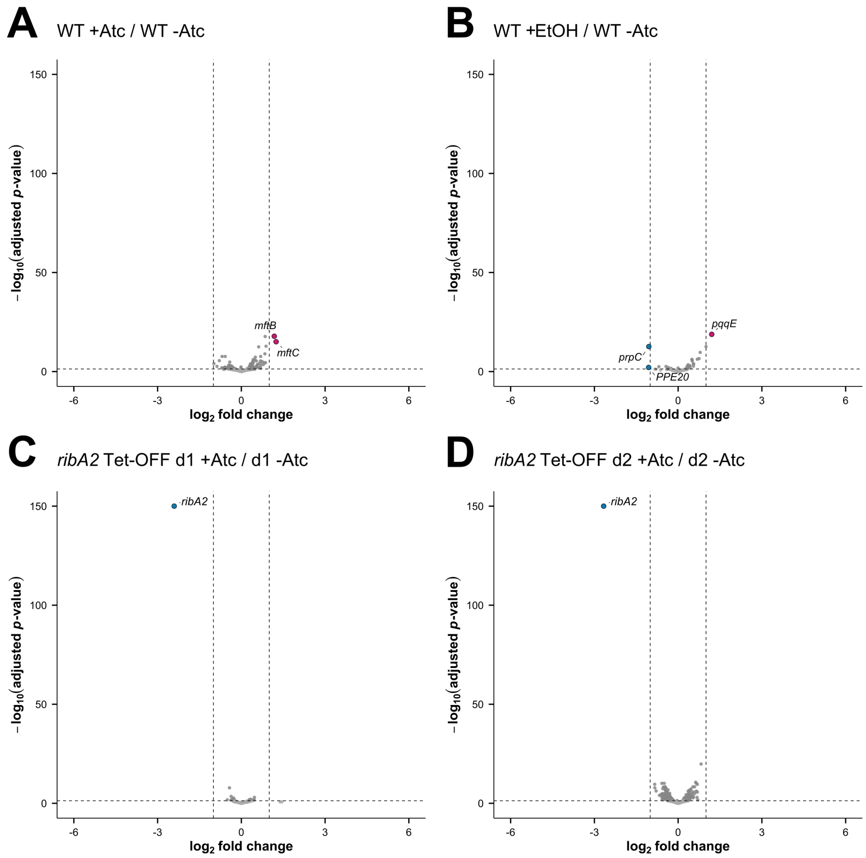


**Figure S2. Differential gene expression in WT or *ribA2* Tet-OFF *M. tuberculosis* at days 1 and 2.** RNA was extracted from WT Erdman or *ribA2* Tet-OFF grown in 7H9 ± 100 ng/ml Atc or in 7H9 +0.1% ethanol (solvent control) and transcriptional profiles were determined by RNAseq. Volcano plots of $\log_2$-fold change and adjusted *P*-value determined by RNAseq. Each dot represents an individual gene. Dashed lines indicate the $\log_2$-fold change = ± 1 and *Padj* < 0.05 (?) statistical significance cutoffs. Red or blue dots represent significantly up- or down-regulated genes relative to the control, respectively. **A.** WT +Atc vs WT no Atc control, day 1. **B.** WT +ethanol vs WT no Atc control, day 1. **C.** *ribA2* Tet-OFF +Atc vs *ribA2* Tet-OFF no Atc, day 1. **D.** *ribA2* Tet-OFF +Atc vs *ribA2* Tet-OFF no Atc, day 2.

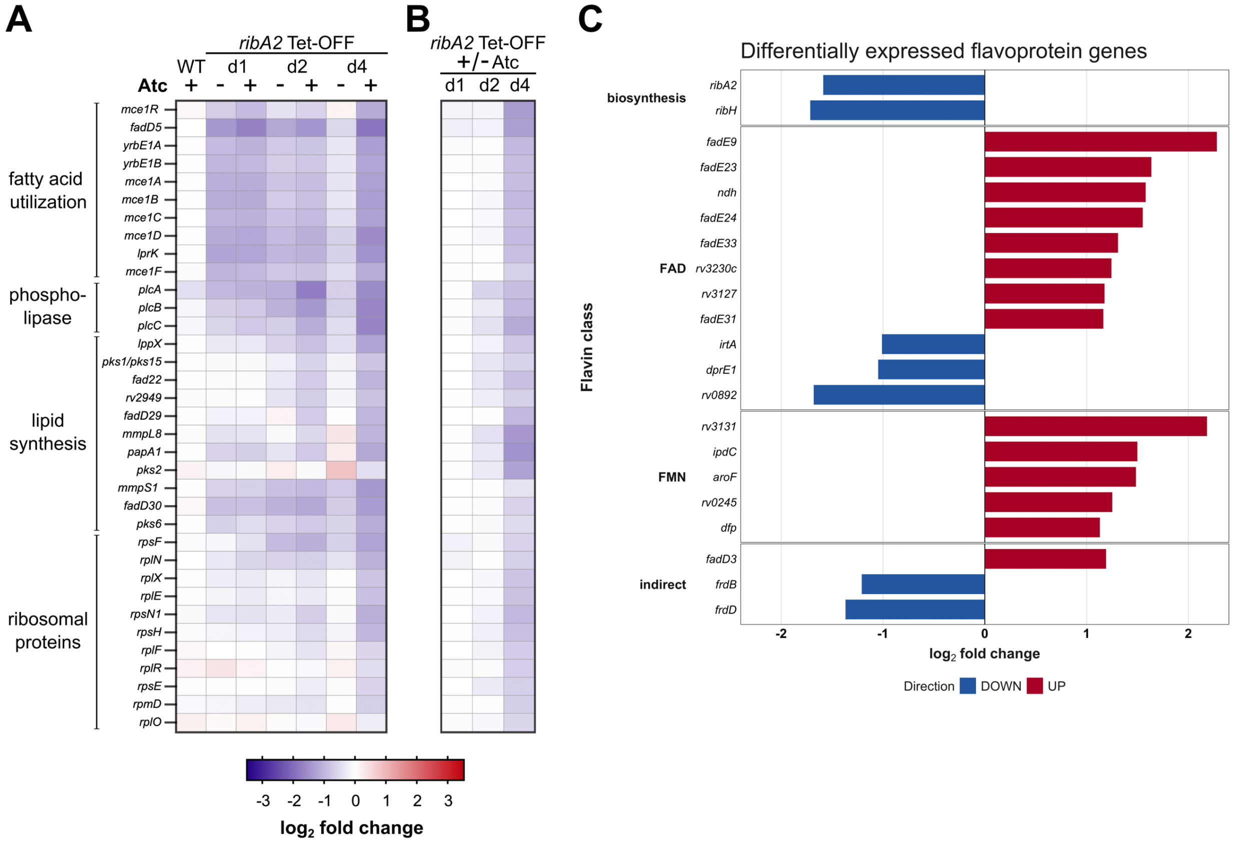


**Figure S3. Lipid metabolism, ribosomal protein genes and flavin-associated genes that are differentially expressed upon *ribA2* transcriptional repression. A-B.** Heat maps showing the $\log_2$-fold change in transcript abundance of differentially expressed genes related to fatty acid utilization, lipid synthesis and ribosomal proteins relative to the WT no Atc control (**A**) or relative to the *ribA2* Tet-OFF no Atc control from the same time point (**B**). **C.** Genes encoding flavoproteins that were significantly differentially expressed in *ribA2* Tet-OFF +Atc day 4 relative to WT no Atc control.

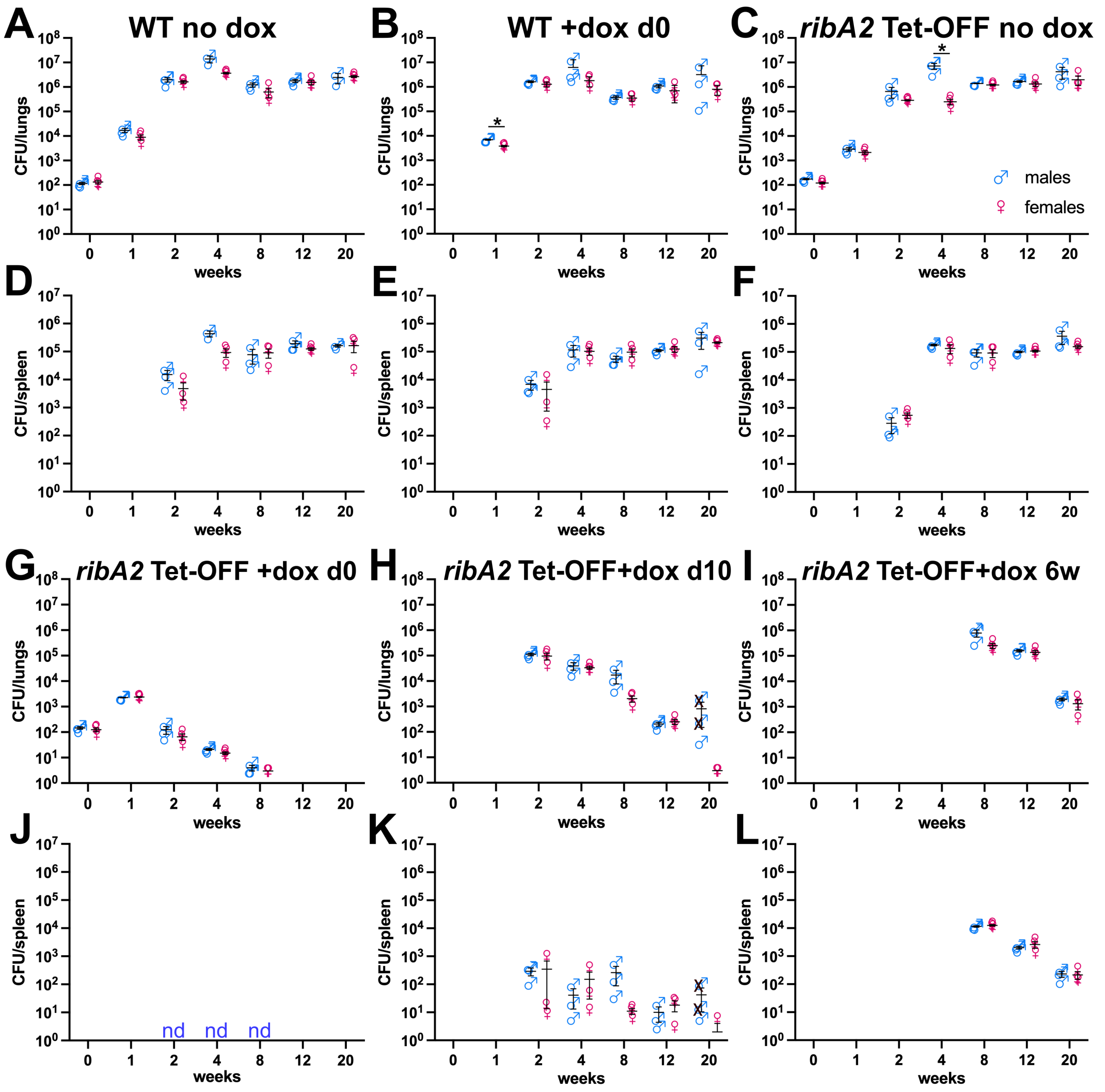


**Figure S4. *M. tuberculosis* lung and spleen burdens from individual C57BL/6 mice, separated by sex.** Data from C57BL/6 mice that were infected by aerosol with WT Erdman (**A, B, D, E**) or *ribA2* Tet-OFF (**C, F, G-L**) and treated with doxycycline (dox, 2,000 ppm in chow) starting at the indicated time points, from Fig 5. *M. tuberculosis* CFU in lungs (**A-C, G-I**) and spleen (**D-F, J-L**) were determined by plating. Data from male and female mice (*n*=3 each) are shown separately. Lines indicate the mean. In **J**, nd indicates no CFU detected in any mouse (detection limit = 3 CFU). Asterisks indicate a statistically significant difference in CFU between male and female mice by unpaired *t*-test, *$P<0.05$. In **H** and **K**, X's over the data for two males indicate mice that were excluded from the averaged data in Fig 5 because *ribA2* Tet-OFF isolates recovered from these mice harbored mutations in *tetR38* (see **Figure S5**).

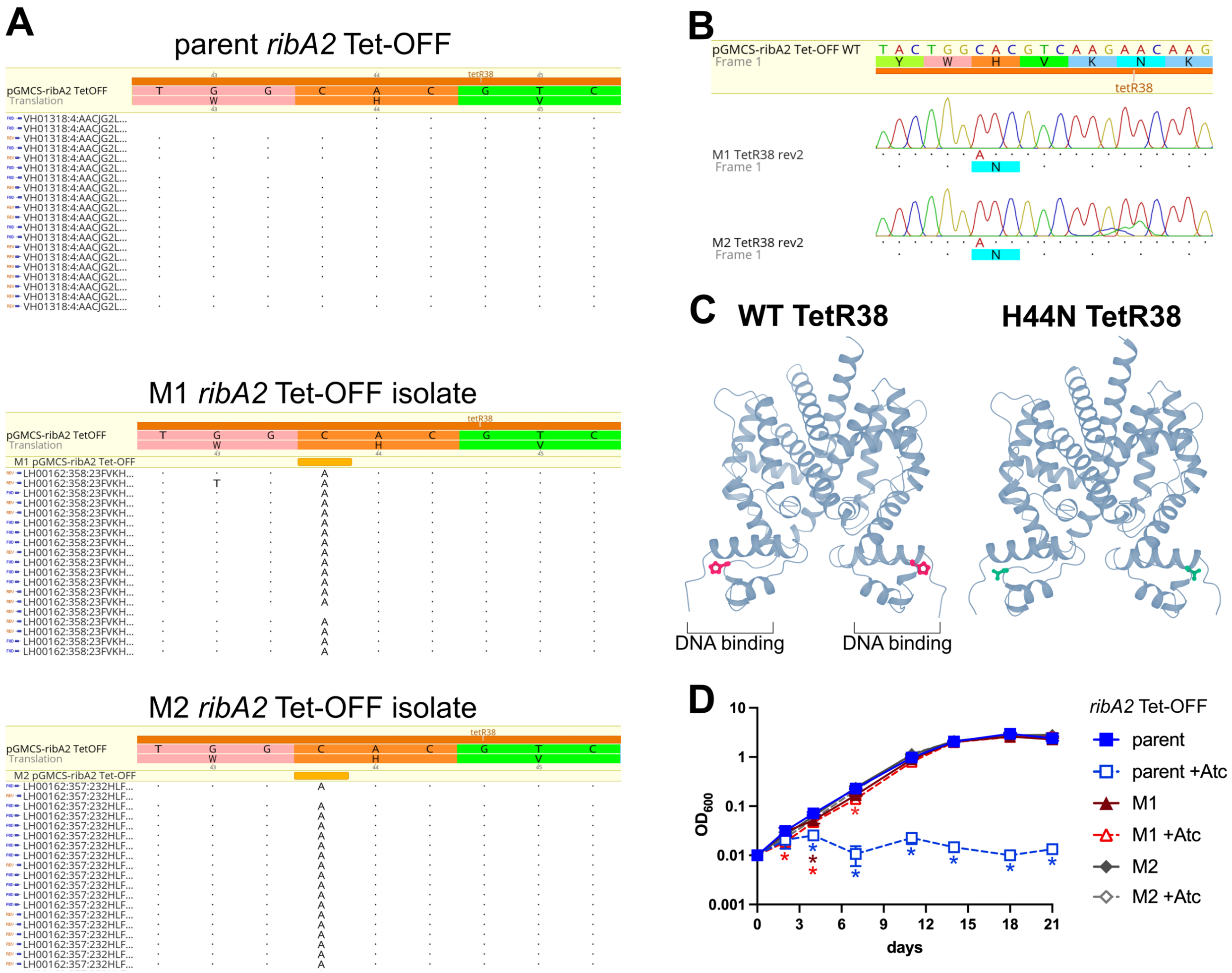


**Figure S5. Mutations in *tetR38* prevent conditional repression of *ribA2* in *ribA2* Tet-OFF isolates from two C57BL/6 mice. A.** Whole genome sequencing reads for the *ribA2* Tet-OFF parental strain and two mouse isolates indicating a consensus C to A single nucleotide substitution in codon 44 of the *tetR38* on pGMCS-*ribA2* in both mouse isolates. **B.** Sanger sequencing confirmation of the A substitution in codon 44 of *tetR38* in both mouse isolates. **C.** Ribbon diagrams of TetR38 (left) and TetR38-H44N (right) protein structures, with the amino acid substitution in the DNA binding domain highlighted. **D.** Growth in complete 7H9 without riboflavin. The *ribA2* Tet-OFF parent strain and *ribA2* Tet-OFF mouse isolates (M1, M2) were grown in complete 7H9 to mid-log phase, then diluted to $OD_{600}$ = 0.01 in 7H9 ± 100 ng/ml anhydrotetracycline (Atc). Growth was monitored by $OD_{600}$ measurement. Data are means ± standard deviations of triplicate cultures. Asterisks indicate statistically significant differences compared to the *ribA2* Tet-OFF parental no Atc control by one-way ANOVA with Dunnett's multiple comparisons test: *$P$<0.05.

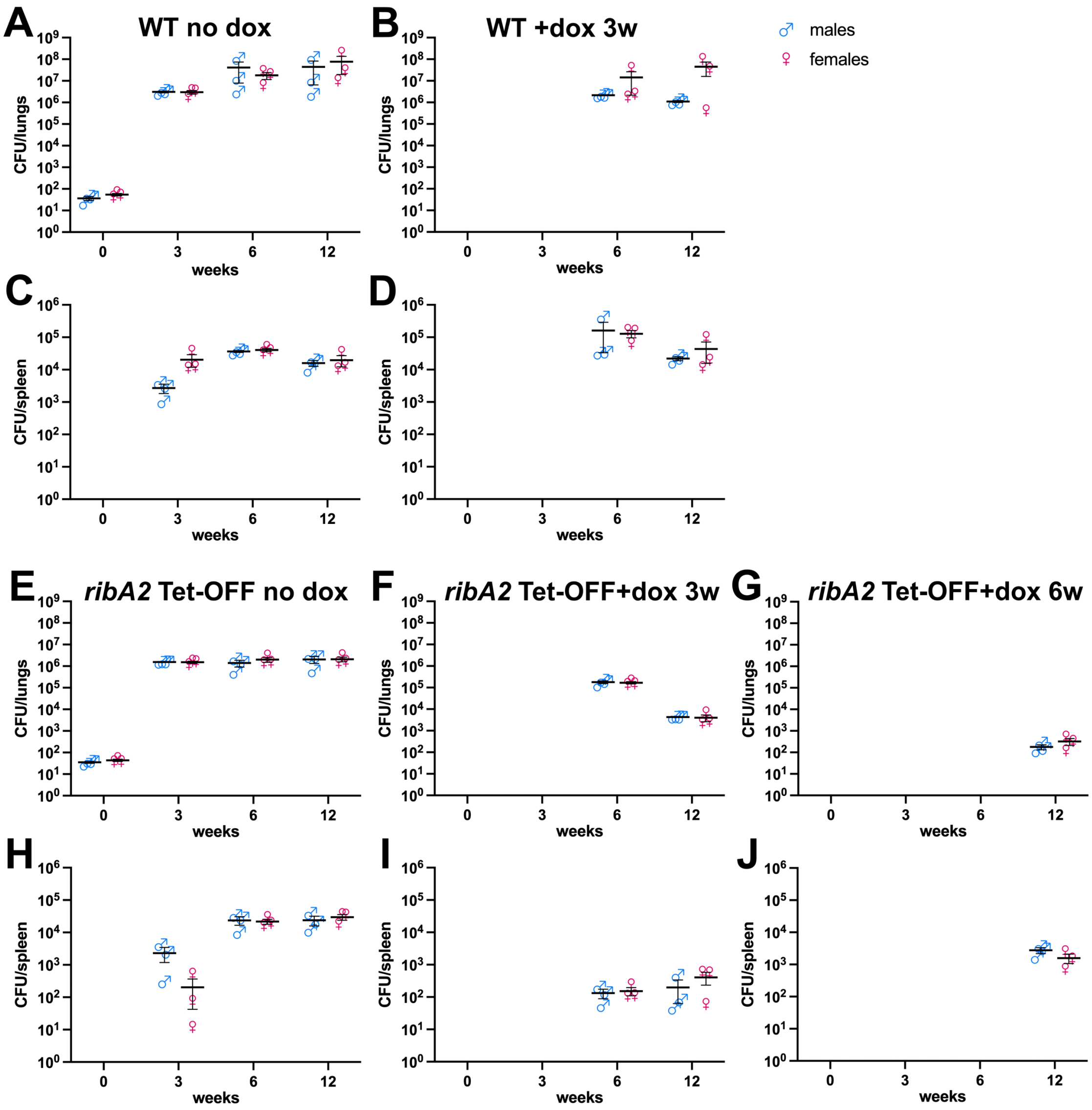


**Figure S6. *M. tuberculosis* lung and spleen burdens from C3HeB/FeJ mice, separated by sex.** Data from C3HeB/FeJ mice that were infected by aerosol with WT Erdman (**A-D**) or *ribA2* Tet-OFF (**E-J**) and treated with doxycycline (dox, 2,000 ppm in chow) starting at the indicated time points, from Fig 6. *M. tuberculosis* CFU in lungs (**A, B, E-G**) and spleen (**C, D, H-J**) were determined by plating. Data from male and female mice ($n$=3 each) are shown separately. Lines indicate the mean. No statistically significant differences between male and female mice were identified by unpaired $t$-test.
